# Extensive mosaicism by somatic L1 retrotransposition in normal human cells

**DOI:** 10.1101/2022.05.18.492429

**Authors:** Chang Hyun Nam, Jeonghwan Youk, Jeong Yeon Kim, Joonoh Lim, Jung Woo Park, Soo A Oh, Hyun Jung Lee, Ji Won Park, Seung-Yong Jeong, Dong-Sung Lee, Ji Won Oh, Jinju Han, Junehawk Lee, Hyun Woo Kwon, Min Jung Kim, Young Seok Ju

**Affiliations:** Graduate School of Medical Science and Engineering (GSMSE), Korea Advanced Institute of Science and Technology (KAIST), Daejeon 34141, Republic of Korea; Genome Insight Inc., Daejeon 34051, Republic of Korea; Department of Internal Medicine, Seoul National University Hospital, Seoul 03080, Republic of Korea; Korea Institute of Science and Technology Information, Daejeon 34141, Republic of Korea; Department of Surgery, Seoul National University College of Medicine, Seoul 03080, Republic of Korea; Department of Life Science, University of Seoul, Seoul 02592, Republic of Korea; Department of Anatomy, School of Medicine, Kyungpook National University, Daegu 41942, Republic of Korea; Department of Anatomy, Yonsei University College of Medicine, Seoul 03722, Republic of Korea; Department of Nuclear Medicine, Korea University College of Medicine, Seoul 02841, Republic of Korea

**Author notes:** Co-corresponding authors **Corresponding authors Young Seok Ju MD PhD**, Associate Professor, Korea Advanced Institute of Science and Technology, Daejeon 34141, Republic of Korea, and Chief Executive Officer, Genome Insight Inc., Daejeon 34051, Republic of Korea, Min Jung Kim MD PhD, Associate Professor, Department of Surgery, Seoul National University College of Medicine, Seoul 03080, Republic of Korea, Hyun Woo Kwon MD PhD, Associate Professor, Department of Nuclear Medicine, Korea University College of Medicine, Seoul 02841, Republic of Korea. Co-first authors with equal contribution (Chang Hyun Nam, Jeonghwan Youk).

## Abstract

Over the course of an individual’s lifetime, genomic alterations accumulate in somatic cells. However, the mutational landscape by retrotranspositions of long interspersed nuclear element-1 (**L1**), a widespread mobile element in the human genome, is poorly understood in normal cells. Here, we explored the whole-genome sequences of 892 single-cell clones established from various tissues collected from 28 individuals. Remarkably, 88% of colorectal epithelial cells acquired somatic L1 retrotranspositions (**soL1Rs**), carrying ∼3 events per cell on average with substantial intra- and inter-individual variances, which was accelerated at least 10-fold during tumourigenesis. Breakpoints of soL1Rs suggested that a few variant mechanisms can be involved in the L1 retrotransposition processes. Fingerprinting of donor L1s using source-specific unique sequences revealed 34 hot L1s, 44% of which were newly discovered in this study, and many ultra-rare hot L1s in the human population showed higher retrotransposition potential in somatic lineages than common sources. Multi-dimensional analysis of soL1Rs with early embryonic developmental relationships, genome-wide methylation, and gene expression profiles of the clones demonstrated that (1) soL1Rs occur from early embryogenesis at a substantial rate, (2) epigenetic activation of hot L1s is stochastically acquired during the wave of early global epigenomic reprogramming, rather than by the sporadic loss-of-methylation at the late stage, and (3) most L1 transcripts in the cytoplasm do not generate soL1Rs in somatic lineages. In summary, this study provides insights into the retrotransposition dynamics of L1s in the human genome and the resultant somatic mosaicism in normal human cells.

## Introduction

Somatic mutations spontaneously accumulate in normal cells throughout an individual’s lifetime from the first cell division^1,2^. Studies have revealed the landscape of the resultant somatic mosaicism in various normal tissues, including the skin, gut, brain, endometrium, blood, embryo, and germline tissues^3-9^. The acquisition of somatic mutations is caused by diverse mutagenic processes, including both endogenous and exogenous mechanisms^10^. These include spontaneous 5-methylcytosine deamination, errors in the DNA replication process, and exposure to mutagens such as tobacco smoking and ultraviolet light^10^. Therefore, studying these mutations provides insights into the characteristics of DNA damage and repair processes in normal human cells.

Previous studies on somatic mosaicism have primarily focused on small nucleotide variants, such as single nucleotide variants (**SNVs**) and indels^3-9^. More complex structural variants, such as genomic rearrangements and mobilization of transposable elements, remain less explored due, in part, to the technical challenges in their accurate detection^11^, particularly at single-cell resolution^12^. In addition, their innate lower burden per cell compared to somatic SNVs requires more systematic and large-scale studies to reveal the overall landscape of structural variants in somatic cells.

L1 retrotransposons are widespread repetitive elements in the human genome and represent approximately 17% of the entire human genome^13^. Evolutionally, L1 retrotransposons are a remarkably successful parasitic unit in the germline, through ‘copying-and-pasting’ themselves at new genomic sites by hitchhiking cellular transcription and translation machinery^14^. However, most of the approximately 500,000 L1s in the human reference genome are now molecular fossils, or unable to transpose further, because they are truncated and have lost their functional potential. Approximately 120 L1s are known as retrotransposition-competent, called “hot L1s”^15,16^. Occasionally, L1 retrotranspositions have been found in the genetic analysis of tissues in several diseases^17-19^, implying their role in the development of human diseases and necessitating a more systematic characterization.

Somatic L1 retrotransposition (**soL1R**) has been extensively explored in cancer tissues^15,20,21^ Compared to other tissues, soL1Rs in cancers are easier to detect because cancer cells are clonally expanded and events are shared by many cells. SoL1Rs are frequently found in specific types of cancers, such as esophageal and colorectal adenocarcinomas^15^, which often lead to disruption of tumour suppressor genes through inducing L1-mediated complex genomic rearrangements^15^. SoL1R detection in non-neoplastic normal cells is technically more challenging because most events are scattered in single cells. Several techniques, such as quantitative polymerase chain reaction (qPCR), L1 reporter assays, L1 captures, and whole-genome amplification, have been employed to reveal soL1Rs in normal cells, such as neurons^22-27^. However, these efforts could not comprehensively and precisely explore single-genomes and reported inconsistent soL1R rates (ranging from 0.04 to 13.7 soL1Rs per neuron). Therefore, the landscape of L1 retrotransposition has not yet been clearly defined in non-neoplastic human cells.

To explore L1 mobilization in healthy human cells, we extensively investigated whole-genome sequences of colonies expanded from single cells collected from human adult tissues (hereafter referred to as ‘clones’). Our approaches allowed for the sensitive and precise investigation soL1Rs in single genomes with minimal amplification and dropout artifacts that frequently occur in whole-genome amplification (WGA)-based single-genome sequencing^28^ and in laser capture microdissection (LCM)-based clonal patch sequencing (**Supplementary Discussion**). In addition, our approach enabled us to combine DNA methylation and gene expression profiles from the same clones, which are challenging in other approaches^29^. Furthermore, by integrating early developmental lineages of clones reconstructed by somatic mutations as cellular barcodes^8^, the early molecular dynamics of the mutational, transcriptional, and epigenetic profiles of L1s were also investigated.

### SoL1R rates in normal colorectal epithelium

To detect soL1R events, we explored 911 whole-genome sequences, including clones from healthy tissues (n=880; 27 individuals), adenomatous polyps (n=12; from four polyps of one patient with MUTYH-associated polyposis) and the matched cancer tissues (n=19; 19 individuals; **Fig. 1a**). The healthy clones consisted of colorectal epithelial cells (406 clones from 19 individuals), fibroblasts from various anatomical locations across the whole body (334 clones from 7 individuals)^8^, and haematopoietic stem and progenitor cells (140 clones from 1 individual)^6^ (**Supplementary Table 1**). Nineteen matched colorectal cancer tissues were acquired from the same individuals from whom the colorectal clones were collected. From these sequences, we assessed the somatically acquired mutations including SNVs, indels, genomic rearrangements, and soL1Rs (**Supplementary Table 1**).

**Figure 1.**
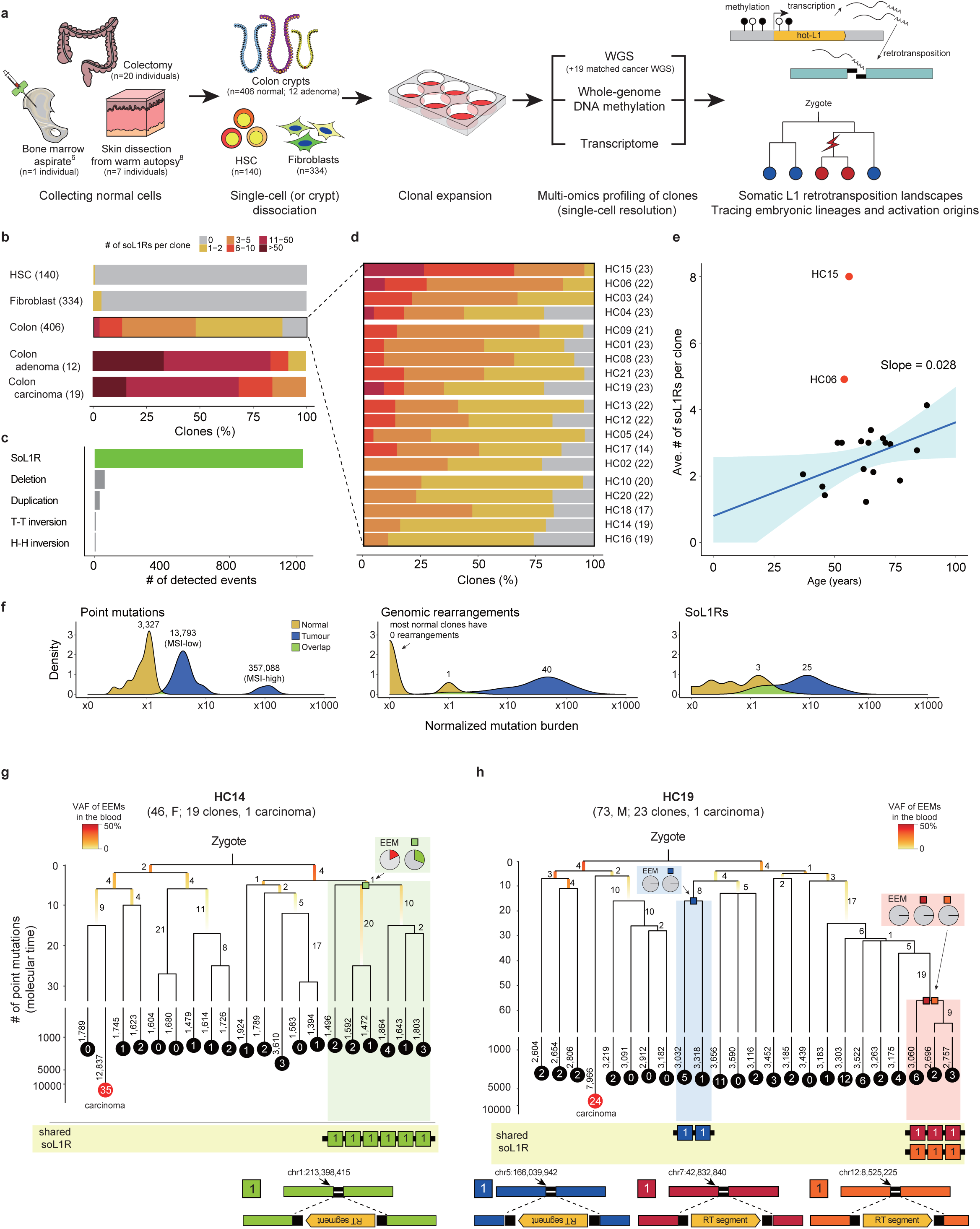
Somatic L1 retrotranspositions in normal cells. **a**, Experimental design of the study. HSC, haematopoietic stem and progenitor cells. WGS, whole-genome sequencing. **b**, Proportion of clones with various numbers of soL1Rs across different cell types. The number of clones for each cell type is shown in parentheses. soL1R, somatic L1 retrotransposition. **c**, Number of structural variations in 406 clones from normal colon epithelial cells. T-T inversion, tail-to-tail inversion; H-H inversion, head-to-head inversion. **d**, Proportion of normal colorectal clones with various numbers of soL1Rs across 19 individuals. The number of clones for each individual is shown in parentheses. **e**, Linear regression of average number of soL1Rs per clone on age. Blue line represents the regression line and shaded areas indicate 95% confidence interval of the regression line. Two outlier individuals (HC15 and HC06) are highlighted in red. **f**, Normalized number of somatic mutations in normal colorectal clones and colorectal cancers. MSI, microsatellite instability. **g, h**, Early phylogenies of normal clones and the matched cancer of HC14 (g) and HC19 (h). A few soL1R events are shared by many clones, suggesting early embryonic retrotranspositions. Branch lengths are proportional to molecular time measured by the number of somatic point mutations shown on the vertical axes. Early branches are coloured by VAFs of early embryonic mutations in the blood. The tips of branches represent normal clone (black dot) or major clone of cancer (red dot), in which the number of soL1Rs is depicted. Pie charts indicate the proportion of blood cells harboring the soL1R. VAF, variant allele fraction; EEM, early embryonic mutation.

Based on the variant allele fractions (**VAFs**) of somatic point mutations, we confirmed that the vast majority of the clones were established from a single founder cell (**Extended Data Fig. 1a**). In addition, genome-wide sequencing depth indicated that the genomic copy number was stable during the clonal expansion, which is expected for non-neoplastic normal cells (**Extended Data Fig. 1b**). Furthermore, somatic mutation burden and no remarkable cancer driver mutations in clones confirmed that these clones were established from non-neoplastic cells (**Supplementary Table 1**).

In the 880 normal clones, we identified 1,250 soL1Rs using a combined analysis of four different bioinformatics tools (**Supplementary Tables 1, 2**). Four lines of evidence indicated that most of the detected soL1Rs were true somatic events that accumulated in vivo rather than culture-induced events. First, the VAFs for soL1Rs were distributed approximately 50%, thus were shared by all cells in a clone (**Extended Data Fig. 1c**). Similarly, in clones established from male donors, soL1Rs in non-pseudo-autosomal regions of chromosome X exhibited approximately 100% VAF. Second, we experimentally confirmed the rate of culture-associated events in 13 pairs of serial single-cell expansions, directly suggesting that >90% of the detected soL1Rs were true in vivo events (**Extended Data Fig. 1d**). Third, we observed a high level of cell-type specificity in the soL1R burden. Fourth, we found a positive correlation between the soL1R burden and the age of individuals. The third and fourth features are not expected if most soL1Rs were acquired by culture-associated artifacts.

As briefly mentioned above, we found a high level of cell-type specificity in the frequency of soL1Rs. For example, 88% of the normal colorectal clones harbored at least one soL1R event (**Fig. 1b**). Remarkably, soL1Rs were more abundant than other classical structural variations in these cells (**Fig. 1c**). In contrast, in fibroblast and blood clones, soL1Rs were mostly absent (P=3.4×10^−171^, two-sided Fisher exact test). On average, we detected approximately three soL1Rs per normal colorectal clone (1,236 soL1Rs in the 406 clones). However, there were substantial variations in soL1R burden within and between individuals. For example, in HC15, the soL1R burden ranged between 1-18 across the 23 clones of the individual (**Fig. 1d**). Overall, 184 soL1Rs were identified from HC15 (8 soL1Rs per clone), which was a 2.6-fold higher number than random expectation (P=9.7×10^−30^, two-sided exact Poisson test).

For each individual, the average soL1Rs rates showed a positive relationship with age, with approximately 0.028 soL1Rs per clone per year (**Fig. 1e**), similar to the clock-like property of endogenous somatic SNVs and indels (**Extended Data Fig. 2a**)^30^. We did not observe strong associations between the rate and the sex and/or anatomical location of the clones in the colon (**Extended Data Figs. 2b, c**). At the individual clone level, no remarkable relationships were observed between the soL1R burden and other somatic mutational statuses, such as point mutation burden, telomere length, the activity of cell-endogenous SNV processes (SBS1 and SBS5; standard signatures in the COSMIC database), exposure to reactive oxygen species (SBS18), and colibactin from *pks*^*+*^ *E. coli*^31^ (SBS88; **Extended Data Figs. 2d-i**), suggesting that soL1R events are not tightly associated with other mutational processes. Collectively, our data indicate that a high level of stochasticity underlies soL1R acquisition at the single-cell level, although the overall chance of soL1R increases broadly over time. Additionally, the higher soL1R burdens in two outlier individuals (HC15 and HC06; **Fig. 1e**) imply a germline predisposition and/or specific environmental exposure that can stimulate L1 retrotransposition.

**Figure 2.**
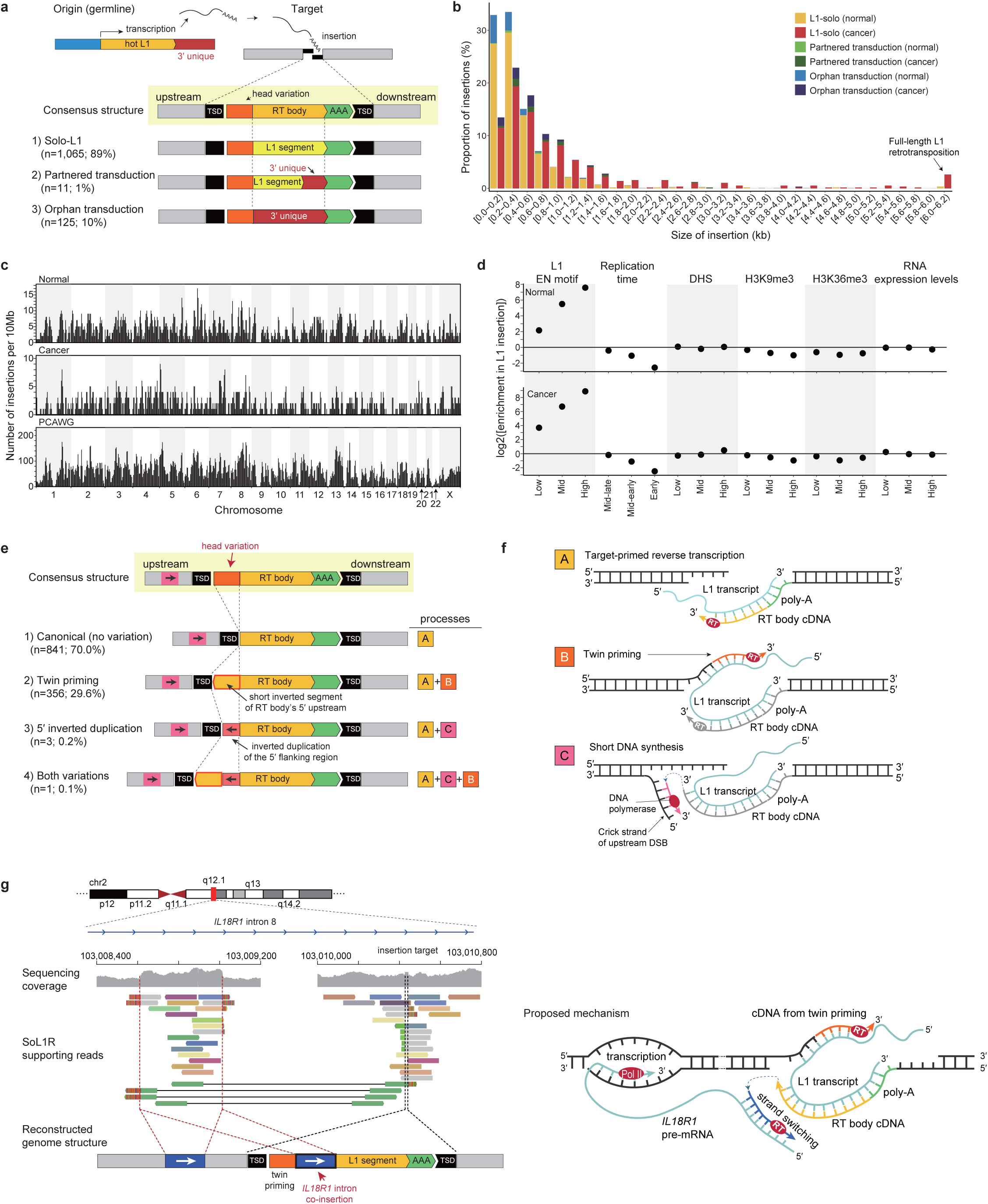
Genomic features of somatic L1 insertion sites. **a**, Schematic diagram of three classes of L1 retrotransposition: solo-L1, partnered transduction, and orphan transduction. RT body, retrotransposed body; TSD, target site duplication. **b**, Distribution of L1 insertion size in normal colorectal clones and colorectal cancers. **c**, Genome-wide distribution of soL1R target sites in normal colorectal clones, colorectal cancers, and the 2,954 pan-cancers analyzed in the PCAWG study (Ref. ^15^). Bars represent the number of L1 insertions in a 10-Mb sliding window with a 5-Mb-sized step. **d**, Association between L1 insertion rate and various genomic features. Dots represent the log value of enrichment scores calculated by comparing bins 1–3 against bin 0 for each feature. L1 EN motif, L1 endonuclease target motif; DHS, DNase I hypersensitivity site. **e, f**, Schematic diagrams of genomic structures of canonical and complex L1 insertions (e) and underlying mechanisms (f). DSB, double-strand break. **g**, An example of a soL1R co-inserted with an expressed gene in the vicinity of the insertion site. A suggestive mechanism is shown in the right panel.

### Acceleration of soL1R rates in tumourigenesis

We compared the soL1R landscape in the normal colorectal clones (1,236 soL1Rs from 406 clones) with those in neoplastic cells, including MUTYH-associated adenomatous polyps (457 soL1Rs from 12 clones) and matched colorectal carcinomas (572 soL1Rs from 19 tissues; **Supplementary Table 2**). All adenoma clones and carcinoma tissues harbored soL1Rs, more frequently than the normal epithelium (100% vs. 88%; *P*= 0.037, two-sided Fisher exact test). The soL1R burden per cell in adenoma and carcinoma was 38 and 30 soL1Rs on average, respectively, with considerable variance, ranging between 2-66 in the 12 adenoma clones and 4-105 in the 19 carcinoma tissues (**Fig. 1b**). The soL1R burden was approximately 10-fold higher in adenomas and carcinomas compared to normal cells, suggesting that the processes of neoplastic transformation provide more favorable conditions for L1 retrotransposition processes. The distribution of soL1R burden showed a larger overlap between normal and cancer than other types of mutations, such as point mutations and genomic rearrangements (**Fig. 1f**).

### L1 retrotransposition starts during embryogenesis

Of the 1,236 soL1Rs detected in the colorectal clones, 30 were redundant (10 events when collapsed), or shared by two or more clones in an individual, implying that these were acquired early in the most recent common ancestral (**MRCA**) cell of the clones. As expected for embryonic events, clones sharing identical soL1R events were the progeny of an ancestral cell in the developmental phylogenies of clones reconstructed by early embryonic mutations (**EEMs**)^8,32^ (**Figs. 1g, h; Extended Data Figs. 3, 4**).

**Figure 3.**
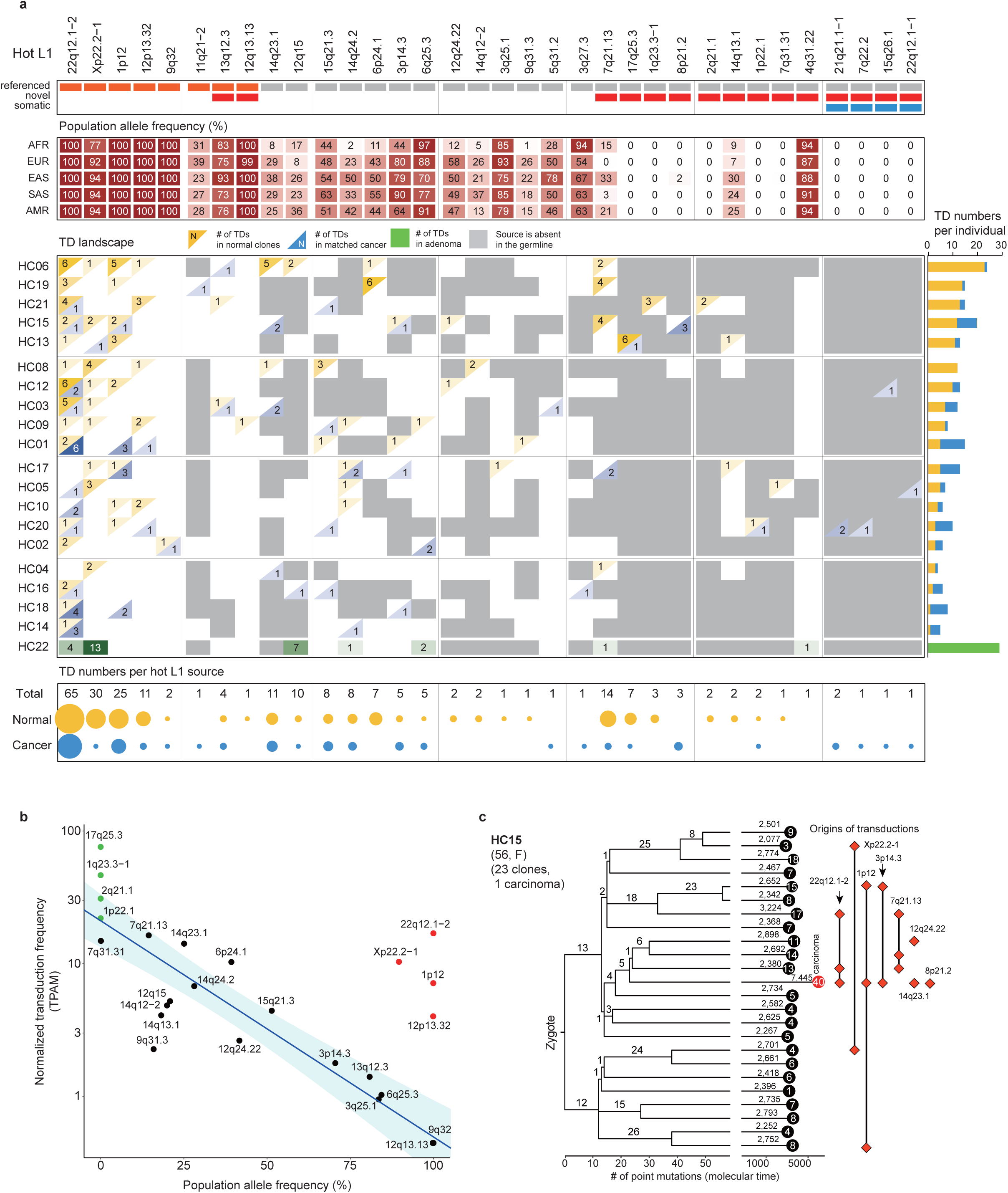
Dynamics of L1 source element activity. **a**, The landscape of transduction events with the features of 34 hot L1s. TD, transduction. **b**, Relationship between the population allele frequency of hot L1 and their normalized retrotransposition activity. Green dots indicate private sources present in only one individual in our colorectal cohort. Red dots indicate common sources, but showing higher retrotransposition activity than expected. TPAM, the number of transductions per allele per 1 million clock-like mutations of molecular time. **c**, Early phylogenies of HC15 clones and the distribution of clones harboring transduction events from specific hot L1 sources. Orange diamonds represent the transduction events from specific hot L1s across colorectal clones. Branch lengths are proportional to molecular time measured by the number of somatic point mutations shown on the horizontal axes. The tips of branches represent normal clone (black dot) or major clone of cancer (red dot), in which the number of soL1Rs is depicted.

**Figure 4.**
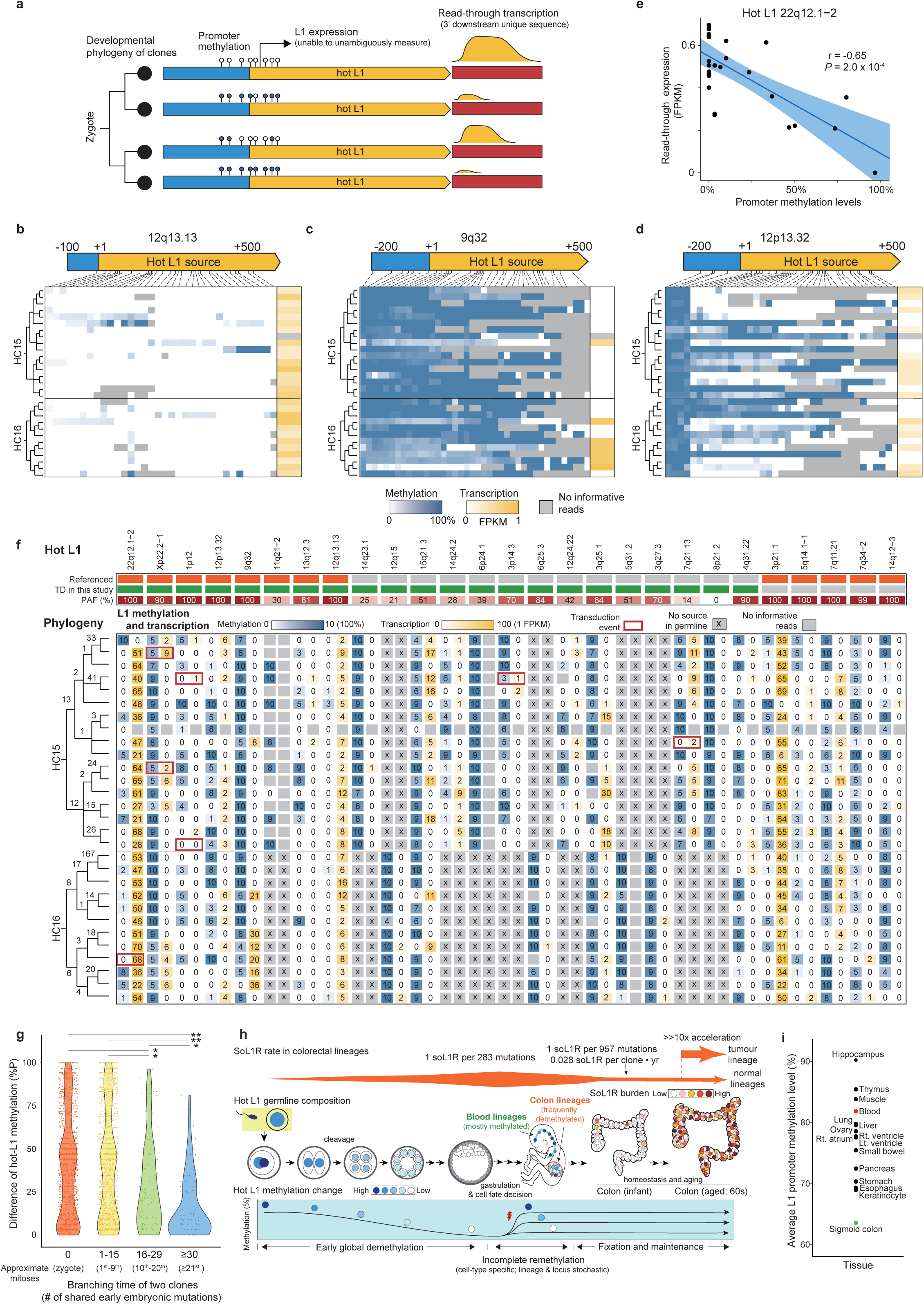
Regulation of L1 source element activity. **a**, Schematic diagram of an integrated analysis of the developmental phylogenies, genome-wide methylation, and gene expression profiles of clones. **b-d**, DNA methylation status in promoter region and read-through RNA expression level of hot L1 at 12q13.13 (b), 9q32 (c), and 12p13.32 (d). FPKM, Fragments per kilobase of transcript per million. **e**, Relationship between DNA methylation status in promoter region and read-through RNA expression level of hot L1 at 22q12.1-2. **f**, A panorama of DNA methylation status, read-through RNA expression levels and developmental phylogenies in 27 hot L1s of 29 normal colorectal clones collected from HC15 and HC16. TD, transduction; PAF, population allele frequency. **g**, Clones branched from early cells after 30 somatic mutations in molecular time exhibit similar hot L1 methylation profiles. Differences of methylation levels of hot L1s in each pair of clones are correlated with the number of early embryonic mutations in their most recent common ancestral cell. Of the 34 hot L1s, 14 showing a substantial methylation variation across clones are selected for the analysis. *P<0.0125, **P<1.25×10^−8^ (Two-sample Kolmogorov-Smirnov test). **h**, Schematic diagram illustrating factors influencing to the soL1R landscape. Composition of hot L1s are genetically determined in the fertilized egg. Demethylation of hot-L1s are acquired during the early development stochastically in each lineage, presumably in the stage of cell fate decisions. SoL1Rs accumulate in the colorectal epithelium throughout life, with ∼0.028 per clone per year. soL1R, somatic L1 retrotransposition. **i**, Average level of L1 promoter methylation in 13 referenced hot L1s across different tissues from ENCODE.

Furthermore, by applying the number of EEMs as a molecular clock (estimated previously in Ref. ^8^: 3.8 mutations per daughter cell for the first two cell divisions in life and 1.2 mutations per daughter cell for the following divisions), the timing of soL1R acquisition during embryogenesis was estimated. For example, a soL1R event in HC14, inserted at chr1:213,398,415 (**Fig. 1g**), was shared by six clones (6 out of 19 clones). Both the common ancestral node in the phylogeny (the second node) and the molecular time (five EEMs) indicated that the event occurred in one embryonic cell at the four-cell stage embryo. Since the four-cell stage embryo stage is before gastrulation, the event was likely shared by multiple germ layers beyond the endoderm (colon). In line with the speculation, we found that the event was shared by 34% of polyclonal blood cells (mesodermal origin; 17% VAF in 199x blood WGS; **Fig. 1g**).

Although the event clearly shows that soL1R is possible at the very early stage embryogenesis, the other nine shared soL1Rs by colorectal clones were absent in blood cells. Their latter node positions in the phylogenies and later molecular time (16-56 EEMs, which is equivalent to the 11^th^-45^th^ cell generations) were consistent with post-gastrulation^8^ (**Fig. 1h; Extended Data Figs. 3, 4**). Collectively with the observation that such embryonic soL1Rs were not found in the 8 phylogenies reconstructed from fibroblasts (7 individuals) and haematopoietic stem and progenitor cells (1 individual), our findings imply that soL1Rs are more substantially activated when embryonic cells are fate determined into colorectal epithelium.

Of 19 phylogenies by normal colorectal clones, informative embryonic lineages (early shared branches) included a total of 2,827 mutations of molecular time (**Extended Data Figs. 3, 4**). Therefore, our observation (10 embryonic soL1Rs) suggests one soL1R per 283 somatic point mutations in the early stage of embryonic development (95% CI: one in every 175-744 mutations; equivalent to 1.6×10^−3^ ∼ 6.8×10^−3^ soL1R per cell per cell division), which was 3.3-fold higher than the rate at the late somatic lineages unique to single clones (one per 940 clock-like mutations in molecular time; P=0.001, two-sided Poisson exact test). Our findings indicated that the soL1R rate is higher in embryogenesis than in somatic lineages after birth.

### Genomic features of soL1R insertions

The genomic consequences of soL1Rs were insertions of L1-related sequences (termed **RT body**), predominantly supported by two imprints of retrotransposition, the poly-A tail and target site duplication (**TSD; Fig. 2a; Extended Data Fig. 5a**). The inserted sequences were most frequently the 3⍰ fragments of the L1 elements (n=1065; 89%; known as solo-L1)^21^. Generally, solo-L1 insertions were restricted to the 3⍰ part of full-L1 sequence (453bp on average); however, full-length mobilizations, which should have the potential for further retrotransposition, were also observed (n=4; 0.4%; **Fig. 2b**).

Additionally, the 3⍰ downstream unique sequence of L1 was often co-inserted with or without the L1 sequence, which are events known as partnered transductions (n=11; 1%) and orphan transductions (n=125; 10%), respectively (**Fig. 2a**)^21^. The relative proportion of transduction among soL1Rs was similar between normal and cancer cells (11.3% vs. 12.4% for normal and cancers, respectively). The average lengths of inserts in the partnered and orphan transductions were 615bp and 245bp on average, respectively. Overall, the length of RT bodies was shorter in normal cells than in colon cancers (453bp vs. 1,031bp for solo-L1, *P*=8.5×10^−20^, two-sided t-test; 615 vs. 755bp for partnered transductions, *P*=0.59, two-sided Wilcoxon rank-sum test; and 245 vs. 530bp for orphan transductions, *P*=0.004, two-sided t-test; **Fig. 2b**), suggesting that polymerization by reverse transcriptase is less processive in normal cells than in cancers.

Although approximately 1% of soL1Rs found in pan-cancers are combined with other genomic rearrangements^15^, we did not find such a complex event in the normal colorectal epithelium, suggesting a high level of negative selection for these changes in non-neoplastic cells. In normal cells, structural variations involving large-scale genome changes would induce cell cycle arrest and negative selection^33^. Similarly, soL1Rs involving exons of protein-coding genes were rare in normal clones, as only one event (0.08%) hit the exonic sequences of a gene (*ASTN1*). The frequency was significantly lower than the random expectation (1 vs. 28; *P*=2.8×10^−11^, two-sided Poisson exact test) and the rate observed in tumours (0.08% vs. 0.52%; *P*=0.026, two-sided Poisson exact test). The lower insertion rate in exons is consistent with the genomic distribution of L1 copies in the germline^34^.

SoL1Rs were distributed across the whole genome (**Fig. 2c**). Some known soL1R hotspots in cancer, including the subtelomeric region of chromosome 5p, were not replicated in normal clones^21^. SoL1Rs in normal clones were more frequently inserted in regions of L1-endonuclease target site motifs (190-fold; 95% confidence interval (CI), 78.9-459) and late replicating regions (5.91-fold; 95% CI, 4.50-7.77; **Fig. 2d**), consistent with previous reports in cancers^15^. Chromatin states and transcriptional levels had a small effect on L1 insertion rates (**Fig. 2d**).

### Insertion processes inferred from the breakpoints

We further investigated breakpoint sequences at the soL1R target sites to infer the mechanistic processes of L1 insertions. In addition to the two canonical features (TSD and poly-A tail), which are acquired by target-primed reverse transcription (**Figs. 2e, f;** by process A), a substantial fraction of soL1Rs showed local sequence deviations, characterized by (1) short inversion in the intra-RT body (n=356; 29.6%), (2) short foldback inversion (inverted duplications) in the 5⍰ upstream of the target site (n=3; 0.2%), or (3) both (n=1; 0.1%; **Fig. 2e**). From the arrangement and orientation of these deviations, we speculate that two basic mechanisms are involved. The former pattern (short inversion in the intra-RT body) is attributable to twin priming^35^, in which the 3⍰ overhang upstream of the double-strand break (DSB) serves as an additional primer for the reverse transcription of L1 transcripts (**Fig. 2f;** by process B). The latter (short foldback inversion) is an imprint of additional DNA synthesis during the final resolution stage of the L1-mediated DSB (**Fig. 2f;** by process C). This pattern suggests that the 3⍰ end of the reverse transcribed sequence is continuing to further DNA replication by DNA polymerase using the Crick strand of the upstream DSB as a template. We observed an additional occasional event in a clone from adenoma, in which a part of the precursor mRNA, transcribed in the vicinity of the insertion site, was reverse-transcribed and co-inserted into the genome, suggesting strand switching of the reverse transcriptase (**Fig. 2g**). These features collectively illustrate that soL1Rs are not acquired by fully ordered linear processes, but many optional events can be engaged stochastically^36^. This may imply functional instability of reverse transcriptase in somatic cells.

Interestingly, we found two clones, each of which had transductions at different genomic target sites but with the identical poly-A tail position in the unique sequences (**Extended Data Fig. 5b**). Given that poly-A tailing is a random event in the transcription of the L1 downstream region, our findings suggest that a single L1 transcript can produce multiple retrotranspositions.

### Hot L1 polymorphism across populations

Fingerprinting of the L1 origin is possible in transductions using the co-inserted 3⍰ unique sequence as L1 barcodes^21^. From the 241 transduction events detected in clones and cancer tissues, 34 hot L1 sources were identified (**Supplementary Table 3**). Of these, 15 (44%) were new and did not overlap with the 124 previously known sources^15^. Of the 15 novel hot L1s, two (13%) were present in both the germline of the clones and the human reference genome (**referenced hot L1s**). Nine (60%) novel hot L1s were present in the germline of the clones but were absent in the reference genome (**non-referenced hot L1s**); therefore, these are polymorphic in human populations. The other four (27%) hot L1s were absent in both the germline and the reference genome, suggesting that the sources were acquired in somatic lineages post-zygotically during their lifetime (**somatically acquired hot L1s; Extended Data Fig. 5c**).

Then, we investigated the population allele frequency (PAF) of 139 hot L1 sources (consisting of 124 known and 15 novel hot L1s; **Supplementary Table 3**) in human populations with a panel of 2,860 individual whole-genomes encompassing five major ethnic groups (714 Africans, 588 Europeans, 538 South Asians, 646 East Asians, and 374 Ad Mixed Americans; **Fig. 3a;** for the full list, **Extended Data Fig. 6**). Although 32 of these were universal in humans (>95% PAF in all ethnic groups), the other 107 (77%) hot L1s were polymorphic, and 37 showed substantial PAF differences (>30%P) across ethnic groups. Intriguingly, 15 hot L1s showed significantly lower PAF in the African population than in non-African populations, suggesting their emergence after the human migration out of Africa. Of note, four hot L1s (17q25.3, 1q23.3-1, 2q21.1, and 1p22.1) were private to an individual in our cohort but not observed in the population panel, indicating that these are ultra-rare sources that are likely to be acquired recently in the germline (**Fig. 3a**). Our observations indirectly suggest that new hot L1 sources are continuously emerging in the human germline, as previously reported^37^, and require more systematic studies to catalog more hot L1s in the pool of the human genome.

### Differential transduction activity of hot L1s

Each of the 34 hot L1 sources contributed to a different number of transduction events across the 20 individuals in the colorectal cohort (**Fig. 3a**). For example, four hot L1s (22q12.1-2, Xp22.2-1, 1p12, and 12p13.32) affected a large fraction (≥ 50%) of individuals and contributed to approximately 50% of the somatic transduction events detected in our study. In particular, 22q12.1-2 caused retrotransposition events in 90% of the individuals and gave rise to more than one-fourth of the transduction events. The high mobilization potential of the source is consistent with previous observations in cancers^15,21^. These four “dominant” hot L1s were universally found in the germline of our cohort (20 out of 20 individuals) and human populations (**Fig. 3a**).

However, the high PAF of hot L1s does not ensure a high mobilization activity. For instance, two hot L1s (9q32 and 12q13.13; 100% PAF) contributed only to the transduction of a single clone while several other known universal hot L1s (n=26) did not generate any transduction events in clones.

To evaluate the normalized retrotransposition activity of hot L1 sources regardless of their abundance in the population, we estimated the absolute transduction rate by counting the number of transductions per L1 allele per 1 million clock-like mutations of molecular time (**TPAM**) in normal clones. Intriguingly, the normalized activity of the sources showed a negative correlation with PAF (**Fig. 3b**). Ultra-rare sources (17q25.3, 1q23.3-1, 2q21.1, and 1p22.1) showed higher TPAM values. In contrast, common hot L1s represented reduced retrotransposition activities, excluding the four dominant hot L1s (22q12.1-2, Xp22.2-1, 1p12, and 12p13.32). These features are consistent with the inverse relationship between prevalence and penetrance of human genomic variants^38^. Because hot L1s cause random insertional mutagenesis, these activities should be repressed in human cells by genetic and epigenetic mechanisms. Ultra-rare hot L1s may maintain their high activity because they emerged more recently, prior to the sufficient negative regulation. It is unclear how the four dominant hot L1s (22q12.1-2, Xp22.2-1, 1p12, and 12p13.32) escaped negative regulation and retained their mobilization potential. We speculate that they either have crucial functional roles or are located in genomic loci that are difficult to repress.

### Epigenetic regulation of hot L1 activation

We then investigated the distribution of embryonic lineages carrying transductions from a specific hot L1 source. There were 31 instances that a hot L1 caused transductions in multiple clones of a single individual (Fig. 3c; **Extended Data Figs. 3, 4**). These clones frequently coalesced in the first or second nodes of the developmental phylogenies. Our findings indicate two different scenarios for the regulation of hot L1: (1) hot L1s are transcriptionally activated in the earliest two- or four-cell stage embryos, or (2) hot L1s are independently switched on in multiple somatic lineages.

To explore the epigenetic and transcriptional dynamics of hot L1s, we integrated genome-wide DNA methylation and RNA expression profiles of clones with the transduction landscapes in clonal phylogenies (**Fig. 4a**). Of note, DNA methylation of the hot L1 promoter has been accepted as a key mechanism for inhibiting L1 transcriptional activation^21,39^. Transcription of each hot L1 source can be specifically assessed by its read-through transcripts in the 3⍰ downstream region^40^.

The profiles of promoter methylation and RNA expression varied across hot L1s (**Figs. 4b-f; Extended Data Fig. 7**). For instance, a hot L1 12q13.13 showed full demethylation for both alleles in most clones, and RNA transcription was generally observed in the 3⍰ downstream region (**Fig. 4b**). In contrast, a hot L1 9q32 represented overall methylation and silent gene expression, particularly in clones from HC15 (**Fig. 4c**). A hot L1 12p13.32 displayed a mixed methylation pattern across clones, predominantly with full methylation and full demethylation (**Fig. 4d**). The L1 promoter methylation profiles showed a strong negative correlation with the RNA expression levels (**Figs. 4e, f**)

The integration of genomic, epigenomic, and transcriptional profiles with the developmental relationship of clones provides three insights into the epigenetic regulation of hot L1s and subsequent genomic insertion.

First, promoter demethylation of hot L1s and its resultant RNA expression are necessary conditions for soL1Rs. Hot L1s that cause transduction events were always demethylated and mostly transcribed in the clone (7 out of 7 instances; highlighted by red rectangles; **Fig. 4f**). However, the opposite was not always true because many clones with demethylated hot L1s did not acquire any transduction events from the source. Our observations also indicated that the demethylation status of hot L1 sources is stable over time. The observed pattern in which active sources are always demethylated is not possible if a hot L1 source is frequently remethylated in somatic lineages after generating a transduction event.

Second, L1 promoter demethylation is driven by the wave of early global epigenetic reprogramming in human embryogenesis rather than erroneous local and stochastic loss-of-methylation in aged cells^41-43^. In clones of epigenetically activated autosomal hot L1s, promoters were frequently fully demethylated for both alleles (**Figs. 4b, d, f**), which was not expected in the latter scenario. Indeed, clones branched from common ancestral cells that existed in the 16-192 embryonic mutations of molecular time in the phylogenetic trees (10^th^-158^th^ cell generations; a common ancestral cell near gastrulation to organogenesis; the reference timing of human embryogenesis in molecular time is available in Ref. ^8^) exhibited similar promoter methylation patterns for a specific hot L1 (**Figs. 4d, f, g; Extended Data Fig. 7**). Our observations are compatible with a scenario in which the L1 promoter is fully demethylated in the earliest global demethylation phase (from the fertilized egg to the blastocyst embryo)^41^ but is not always completely remethylated in the following organogenesis period, particularly in cell lineages differentiating to the colorectal epithelium (**Fig. 4h**). The promoter epigenotypes are then stably inherited through downstream mitoses, like X-inactivation^46^. Given the low soL1R rates in the blood and fibroblasts, we speculate that such incomplete L1 promoter remethylation should be less frequent in embryonic lineages differentiating into those cell types. In line with our speculation, the colon tissues showed the lowest promoter methylation level for hot L1s^44,45^ (**Fig. 4i**). However, it is unclear why embryonic lineages to a specific cell type (i.e., colorectal epithelium) and particular hot L1s (i.e., 12q13.13; **Figs. 4b, f**) are more resistant to the promoter remethylation^47^.

Third, most L1 transcripts are infertile regarding the retrotransposition events in normal cells. Our data suggest that somatic cells, particularly the colorectal epithelium, are continuously exposed to hot L1 transcripts in the cytoplasm. The overall transcription level of hot L1s is approximately 0.5-2 FPKM (**Fig. 4f; Extended Data Fig. 7**) in most colorectal colonies. At face value, it is approximately one hot L1 transcript per cell^48^, but there would be higher number of hot L1 transcripts in the cytoplasm as our L1 expression levels were estimated from 3⍰ read-through transcripts only. Despite the exposure of genomic DNA to L1 transcripts for thousands of cell divisions over lifetime, only three soL1Rs are to be acquired per somatic lineage. Our findings indicate that a series of post-transcriptional processes of L1, including translation of L1 open reading frames, nuclear import of L1 riboprotein, formation of DSBs at target sites, reverse transcription of L1 transcript, and final genomic repair are collectively inefficient. Acceleration of L1 insertion in tumours (**Fig. 1b**) implies that a defense mechanism is more actively operative in normal cells.

## Discussion

Our findings demonstrated that cell-endogenous L1 elements lead to retrotransposition in normal somatic lineages and that colon epithelial cells acquire 0.028 soL1R events per year (**Fig. 4i**). Given the number of crypts in the colon (10 million)^49^, individuals in their 60s would have approximately 20 million retrotranspositions collectively in the colorectal epithelium. A fraction of these L1 insertions can confer phenotypic changes in mutant cells and cause human diseases such as cancer^15^.

Despite the small number of individuals enrolled in this study, we detected many novel hot L1s, implying the presence of additional undetected hot L1s in the pool of human genomes. As shown in our study, some rare hot L1s have higher mobilization activity than common sources. For a more comprehensive catalog of hot L1s, genomic studies on diverse ethnic groups will be helpful. Of note, most of the individuals investigated in this study were Koreans, which has not been a major ethnic group in previous genome studies.

Several complementary methods, such as whole-genome amplification^5^, duplex DNA sequencing^51^, laser-capture microdissection^7,9,32,52^, and *in vitro* single-cell expansions^4,6,8^, can be used to explore somatically acquired genomic changes in normal cells. Although clonal expansions are labor-intensive and only applicable to dividing cells, they have fundamental advantages, including (1) implementing the most sensitive and precise mutation detection at the absolute single-cell level, (2) facilitating additional multi-omics profiling in the same single cells, and (3) permitting the exploration of early developmental relationships of single cells. As shown in this study, these advantages are essential for understanding the multidimensional dynamics of L1 retrotransposition.

In this study, we observed a substantial level of somatic mosaicism in normal cells driven by soL1Rs. However, many things are yet to be discovered. For example, some other cell types not investigated in this study may have higher soL1R rates than colorectal epithelium. As in APOBEC-mediated mutagenesis^50^, acquisition of soL1Rs may be episodic, *i*.*e*., a few soL1Rs can occur together at a specific cell cycle. The sequence polymorphisms of a hot L1 source among individuals may be an important factor for understanding differential retrotransposition activity across individuals^53^ although it could not be systematically explored due to the inherent limitations of short-read sequencing on repetitive sequences. Therefore, further studies using similar approaches but with more innovative sequencing techniques, on a larger number of single genomes from additional cell types, collected at various time points in the course of aging and disease progression will elucidate the panorama of L1 retrotransposition in the human body and their functional impact on disorders.

## Supporting information

Supplementary Discussion

## Methods

### Human tissues

For the in vitro establishment of clonal organoids from the colorectal tissues, healthy mucosal tissues were obtained from surgical specimens of 19 patients undergoing elective tumour-removing surgery (**Supplementary Table 1**). Normal tissues approximately 1×1×1 cm3 in size were cut out from a region > 5 cm away from the primary tumour. Matched blood and colorectal tumour tissues from the same patients were also collected.

Fresh biopsies from one patient of MUTYH-associated familial adenomatous polyposis were obtained from the colonoscopy. Tissues approximately 0.5×0.5×0.5 cm3 in size were cut out from the four polyps. Matched blood and buccal mucosa tissue from the same patient were also collected.

All tissues were transported to the laboratory for organoid culture experiments within eight hours of the collection procedure. All the procedures in this study were approved by the Institutional Review Board of Seoul National University Hospital (approval number: 1911-106-1080) and Korea Advanced Institute of Science and Technology (approval number: KH2022-058). We obtained informed consent from all participants. This study was conducted in accordance with the Declaration of Helsinki and its later amendments. No statistical methods were used to predetermine sample size.

### Publicly available datasets

We included publicly available whole-genome sequences of single-cell expanded clones to reach a more complete picture of L1 retrotransposition in various human tissues. We included 474 whole-genome sequences from two previous datasets, one for haematopoietic cells (140 clones from one individual)^6^, and one for mesenchymal fibroblasts from our previous work (334 clones from seven individuals)^8^.

To understand the PAF of hot L1s, we collected 2,852 publicly available whole-genome sequences of normal tissues with known ethnicity information. These data were collected from various studies ^54-59^.

### Organoid culture of colorectal crypts

All organoid establishment procedures and media compositions were adopted from previous literature with slight modifications^60^. Mucosal tissues were cut into approximately 5 mm and washed with PBS. Tissues were transferred to 10 mM EDTA (Invitrogen) in 50 ml conical tubes, followed by shaking incubation for 30 min at room temperature. After incubation, the tubes were gently shaken to separate crypts from the connective tissues. The supernatant was collected and 20 μl of suspension was observed under a stereomicroscope to check the presence of crypts. Crypt suspension was centrifuged at 300 rcf for 3 min, and the pellet was washed one time with PBS to reduce ischemic time. Isolated crypts were embedded in growth-factor reduced (GFR) Matrigel (Corning) and plated in a 12-well plate (TPP). Plating crypts was at a limited dilution by modifying the protocol from previous literature^61^. Briefly, approximately 2,000 crypts were transferred to 900 μl of Matrigel and plate 3×150 μl droplets in 3 wells of a 12-well plate. Next, 450 μl of Matrigel was added to the remaining dilution and plating of 3 droplets in 3 wells was repeated. Serial dilution was performed at least 4 times and the final remaining dilution was plated in 6 wells. The plates were transferred into an incubator at 37 °C for 5-10 min to solidify Matrigel. Each well was overlaid with 1 ml of organoid culture media. Organoid culture media compositions for the colorectal epithelial cells were described in **Supplementary Table 4**.

### Clonal expansion of single crypt-derived organoid

Primary culture of bulk and diluted crypts was maintained for at least 10 days to ensure the initial mass of single crypt-origin organoid. After growing organoids, a single organoid was manually picked up using a 200 μl pipette under an inverted microscope. Picked organoid was placed into an Eppendorf tube and dissociated using a 1cc syringe with a 25 G needle under TrypLE Express (Gibco). Then, blocking TrypLE by ADF+++ (Advanced DMEM/F12 with 10 mM HEPES, 1X Glutamax, and 1% penicillin/streptomycin) was followed by centrifugation and washing. Pellet was placed in a single well of a 24-well plate. Plates were transferred to a humidified 37 °C/5 % CO2 incubator and the medium was changed every 2-3 days. After successful passage, clonal organoids were transferred to a 12-well plate and further expanded. The confluent clones were collected for DNA extraction and organoid stock.

### Re-clonalization of single crypt-derived organoid

Cultured single crypt-derived organoids were harvested and dissociated using TrypLE Express. After blocking TrypLE and washing, organoids were resuspended using ADF+++. Organoid suspensions were filtered through a 40 μm strainer (Falcon). Then single cells were sorted into a FACS tube by cell sorter (FACSMelody, BD Biosciences). Single cells were selected based on forward- and side-scatter characteristics according to the manufacturer’s protocol. Sorted cells were sparsely seeded with GFR matrigel (500 / well) in 12-well plates. Grown re-clonalized single organoids were manually picked and expanded by the methods described above.

### Library preparation and whole genome sequencing

We extracted genomic DNA materials from clonally expanded cells and matched peripheral blood and colorectal tumour tissue using DNeasy Blood and Tissue kits (Qiagen) or Allprep DNA/RNA kits (Qiagen) according to the manufacturer’s protocol. DNA libraries for whole-genome sequencing (WGS) were generated using Truseq DNA PCR-Free Library Prep Kits (Illumina). WGS was performed on either the Illumina HiSeq X Ten platform or the NovaSeq 6000 platform to generate mean coverage of 17.0× for 406 clonally expanded normal crypts, 34.0×12 crypts from adenomatous polyps, 34.6×19 matched tumour and 173× for 20 matched blood tissues.

### Whole transcriptome sequencing of the organoids

Total RNA from cultured clonal organoids was extracted using Allprep DNA/RNA kits (Qiagen). Total RNA sequencing library was constructed using Truseq Stranded Total RNA Gold kit (Illumina) according to the manufacturer’s protocol.

### Whole genome DNA methylation sequencing of organoids

Genomic DNA was extracted from clonally expanded cells using DNeasy Blood and Tissue kits (Qiagen) or Allprep DNA/RNA kits (Qiagen). The libraries were prepared from 200ng of input DNA with control DNA (CpG methylated pUC19 and CpG unmethylated lambda DNA) using NEBNext Enzymatic Methylation-seq kit (NEB) according to the manufacturer’s protocol. Paired-end sequencing was performed using NovaSeq 6000 platform (Illumina).

### Variant calling and filtering of WGS data

Sequenced reads were mapped to the human reference genome (GRCh37) using the BWA-MEM algorithm^62^. The duplicated reads were removed by either Picard (available at http://broadinstitute.github.io/picard) or SAMBLASTER^63^. We identified single-nucleotide variants and short indels as previously reported^8^. Briefly, base substitutions and short indels were called using Haplotypecaller2^64^ and VarScan2^65^. To establish high-confidence variant sets, we removed variants with the following features: 1% or more VAF in the panel of normals, high proportion of indels or clipping (>70%), 3 or more mismatched bases in the variant reads, and frequent existence of error reads in other clones.

### Reconstruction of the early phylogenies

We reconstructed the phylogenetic tree of the colonies and the major clone of the cancer tissue from an individual by generating an n × m matrix representing the genotype of n mutations of m samples as previously conducted^8^. Briefly, single-nucleotide variants and short indels from all samples of an individual were merged. Only variants with 5 or more mapped reads in all samples were included to avoid incorrect genotyping for the low coverage. Additionally, variants with less than 0.25 VAF in all samples were removed to exclude possible sequencing artifacts. If the VAF of ith mutation in the jth sample is more than 0.1, Mij was assigned 1; otherwise, 0. Mutations shared in all samples were regarded as germline variants and discarded. We grouped all mutations according to the types of samples in which they were found and established the hierarchical relationship between mutation groups. In short, if the samples of mutation group A contain all the samples of mutation group B in addition to other samples, mutation group B is subordinate to mutation group A. Then, we reconstructed the phylogenetic tree that can best explain the hierarchy of the mutation groups. The final phylogenetic tree is a rooted tree where each sample (colony) is attached to one terminal node of the tree, with the number of mutations in the corresponding mutation group being the length of the branch. To convert the molecular time (# of early mutations) to physical cell generations, we used a mutation rate of 3.8 per cell per cell division (pcpcd) for the first two cell divisions and then 1.2 pcpcd, which were estimated from the previous work^8^.

### Calling structural variations

We identified somatic structural variations in a similar way to our previous report^8^. We called structural variations using DELLY^66^ with matched blood samples and phylogenetically distant clones to retain both early embryonic and somatic mutations. Then, we discarded variants with the following features: presence in the panel of normals, an insufficient number of supporting read pairs (less than 10 read pairs without supporting SA tag or less than 3 discordant read pairs with 1 supporting SA tag), and many discordant reads in matched blood samples. To remove remaining false-positive events and rescue false-negative events located near the breakpoints, we visually inspected all the rearrangements passed the filtering process using the Integrative Genomics Viewer^67^.

### Calling L1 retrotransposition

We called L1 retrotranspositions using MELT^68^, TraFiC-mem^21^, DELLY^66^, and xTea^69^ with matched blood samples and phylogenetically distant clones to retain both early embryonic and somatic mutations. Potential germline calls, overlapping with events found in the unmatched blood samples, were removed. To confirm the reliability of the calls and remove remaining false-positive events, we visually inspected all the soL1R candidates focusing on two supporting evidence, 1) poly-A tails and 2) target site duplications using the Integrative Genomics Viewer^67^. Additionally, we excluded variants with a low number of supporting reads (lower than 10% of total reads) to exclude possible artifacts. We obtained the 5⍰ and 3⍰ ends of the inserted segment to calculate the size of soL1Rs and to determine whether L1-inversion or L1-mediated transduction was combined. When both ends of the insert were mapped on opposite strands, the variant was considered to be inverted. When the inserted segment was mapped to unique and non-repetitive genomic sequences^21^ where a full-length L1 element is located within a 15-kb upstream region, we determined the L1 insertion was combined with 3⍰ transduction and derived from the L1 element on the upstream region of unique sequences.

### Population allele frequency of L1 sources

To calculate the PAF of hot L1 sources, we collected 2,852 publicly available and 8 in-house (overall 2,860) whole-genome sequences of normal tissues with known ethnicity information (714 Africans, 588 Europeans, 538 South Asians, 646 East Asians, and 374 Ad Mixed Americans). Initially, we determined whether individuals have hot L1s in their genomes or not. Briefly, we calculated the proportion of L1-supporting reads for non-reference L1 and the proportion of reads with small insert size opposing L1 deletion for reference L1, respectively. Only hot L1s with a proportion of 15% or more were considered to exist in the genome. Then, we calculated the PAF of a specific hot L1 as the proportion of individuals with the L1 in the population.

### Mutational signature analysis

To extract mutational signatures in our samples, we used three different tools (in-house script, SigProfiler^70^, and HDP^71^) to achieve a consensus set of mutational signatures for each type of colon sample, including the normal epithelial cells, adenoma, and carcinoma. Briefly, our in-house script is based on non-negative matrix factorization (NMF), with or without various mathematical constraints, and borrows core methods from the predecessor of SigProfiler^72^, such as using a measure of stability and reconstruction error for model selection; however, it provides more flexibility in examining a broader set of possible solutions, including those that can be missed by SigProfiler, and enables a deliberate approach for determining the number of presumed mutational processes. As a result, we selected a subset of signatures that best explain the given mutational spectrum: SBS1, SBS5, SBS18, SBS40, SBS88, SBS89, ID1, ID2, ID5, ID9, ID18, and IDB for the normal colorectal epithelial cells, SBS1, SBS5, SBS18, SBS36, SBS40, ID1, ID2, ID5, ID9 for MUTYH-associated adenoma, and SBS1, SBS2, SBS5, SBS13, SBS15, SBS17a, SBS17b, SBS18, SBS21, SBS36, SBS40, SBS44, SBS88, ID1, ID2, ID5, ID9, ID12, ID14, and ID18 for colorectal cancers. All signatures are attributed to known mutational signatures available from version 3.2 of the COSMIC mutational signature (available at https://cancer.sanger.ac.uk/cosmic/signatures) and IDB, which is a newly found signature from previous research on the normal colorectal epithelial cells^73^ but not yet cataloged in COSMIC mutational signature.

### Association with genome features

The L1 insertion rate was calculated as the total number of soL1Rs per sliding window of 10Mb with an increment of 5 Mb. To examine the relationship between L1 insertion rate and other genomic features at single-nucleotide resolution, we used a statistical approach described in previous literature^15,74^. In brief, we divided the genome into four bins (0-3) for each of the genomic features, including replication time, DNA hypersensitivity, histone mark (H3K9me3 and H3K36me3), RNA expression, and closeness to L1 canonical endonuclease motif (here defined as TTTT|R (where R is A or G) or Y|AAAA (where Y is C or T)). By comparing the breakpoint sequences with L1 endonuclease motif, we assigned the genomics regions with more than four (most dissimilar), three, two, and less than one (most similar) mismatches to L1 endonuclease motif into bins 0, 1, 2, and 3, respectively. DNA hypersensitivity and histone mark data from the Roadmap Epigenomics Consortium were summarized by averaging the fold-enrichment signal across eight cell types. Then, genomic regions with fold enrichment signal lower than 1 belonged to bin 0, while the remainder was divided into three equal-sized bins: bin 1(least enriched), bin 2(moderately enriched), and bin 3(most enriched). RNA-seq data was also obtained from Roadmap and FPKM (Fragments Per Kilobase of transcript per Million) and averaged across eight cell types. Then, regions with no expression (FPKM = 0) belong to bin 0 while the remainder was divided into three equal-sized bins: bin 1 (least expressed), bin 2 (moderately expressed), and bin 3 (most expressed). Replication time was processed by averaging eight ENCODE cell types, and genomic regions were stratified into four equal-sized regions: bin 0 contains regions with the latest replicating time while bin 3 contains regions with the earliest replicating time. Then, we performed negative binomial regression with all genomics features as covariates. For every feature, enrichment scores were calculated by comparing bins 1–3 against bin 0. Therefore, the log value of the enrichment score for bin 0 should be equal to 0 and is not described on plots.

### Methylation analysis

Sequenced reads were processed using Cutadapt^75^ to remove adaptor sequences. Trimmed reads were mapped using Bismark^76^ to the genome combining human reference genome (GRCh37) modified by incorporating L1 consensus sequences at the non-reference L1 source sites, pUC19, and lambda DNA sequences. For a single CpG site, the number of reads supporting methylation (C or G), the number of reads supporting unmethylation (A or T), and the proportion of the former reads among total reads (methylation fraction) were calculated using Bismark^76^. The conversion efficacy was estimated with reads mapped on CpG methylated pUC19 and CpG unmethylated lambda DNA. To take a look at the overall methylation status, we examined the methylation fraction in regions ranging from 600 bp upstream to 600 bp downstream from the L1 transcription start site for each L1 source element. Then, we focused on the CpG sites located from the L1 transcription start site to the 250 bp downstream region (+1 ∼ +250) and classified each CpG site into three categories according to methylation fraction: homozygous unmethylation (methylation fraction < 25%), heterozygous (methylation fraction ≥ 25% and methylation fraction < 75%), and homozygous methylation (methylation fraction ≥ 75%). Next, methylation scores were assigned to CpG sites (0 for homozygous unmethylation, 5 for heterozygous, and 10 for homozygous methylation) and summarized by averaging the score of all CpG sites on +1 ∼ +250 region of L1 element. Finally, we compared the methylation score across every sample and every known source element to figure out the relationship between methylation status and source activation.

For the analysis of L1 promoter methylation level in bulk tissues, we downloaded WGBS data of 16 different tissues from Roadmap Epigenomics. The Roadmap codes are E050 BLD.MOB.CD34.PC.F (Mobilized_CD34_Primary_Cells_Female), E058 SKIN.PEN.FRSK.KER.03 (Penis_Foreskin_Keratinocyte_Primary_Cells_skin03), E066 LIV.ADLT (Adult_Liver), E071 BRN.HIPP.MID (Brain_Hippocampus_Middle), E079 GI.ESO (Esophagus), E094 GI.STMC.GAST (Gastric), E095 HRT.VENT.L (Left_Ventricle), E096 LNG (Lung), E097 OVRY (Ovary), E098 PANC (Pancreas), E100 MUS.PSOAS (Psoas_Muscle), E104 HRT.ATR.R (Right_Atrium), E105 HRT.VNT.R (Right_Ventricle) E106 GI.CLN.SIG (Sigmoid_Colon), E109 GI.S.INT (Small_Intestine), and E112 THYM (Thymus). The methylation fractions of CpG sites in referenced L1 sources were collected and summarized by averaging the fraction of all CpG sites on +1 ∼ +250 region of L1 element. Then, we compared the averaged L1 promoter methylation level across different tissues.

### Gene expression analysis

Sequenced reads were processed using Cutadapt^75^ to remove adaptor sequences. Trimmed reads were mapped to the human reference genome (GRCh37) using the BWA-MEM algorithm^62^. The duplicated reads were removed by SAMBLASTER59. To identify the expression level of each L1 source element, we collected the reads mapped on the regions up to 1kb downstream from the 3⍰ end of the source element and calculated the FPKM value. Only reads in the same direction with the source element were considered. If the source element is located on the gene and both are on the same strand, the FPKM value was not calculated because the origin of reads on the downstream region is ambiguous.

## Data availability

Whole-genome, DNA methylation, and transcriptome sequencing data are deposited in the European Genome-phenome Archive (EGA) with accession EGAS00001006213 and available for general research use.

## Code availability

In-house scripts for analyses are available on GitHub (https://github.com/ju-lab/colon_LINE1)

## Acknowledgements

We thank Seongyeol Park (Genome Insight Inc.) for his fruitful comments and discussions. This work was supported by the National Research Foundation (NRF) of Korea funded by the Korean Government (NRF-2020R1A3B2078973 to Y.S.J.), and Suh Kyungbae Foundation (SUHF-18010082 to Y.S.J.).

## Author contributions

J.Y. and Y.S.J. conceived the study. J.Y., H.W.K., and J.Y.K. developed the entire protocol of the clonal expansion of the colorectal epithelial cells and conducted experiments. H.J.L., J.W.P., S.-Y.J. and M.K. collected colorectal samples and clinical histories of the patients. S.A.O. conducted genome sequencing. C.H.N. and J.Y. conducted most of genome and statistical analyses with a contribution of Jo.L., H.W.K. and Y.S.J.. J.W.P., Ju.L contributed to large-scale genome data management. D.S.L., J.W.O. and J.H. participated in the data interpretation. C.H.N., H.W.K. and Y.S.J. wrote the manuscript with contributions from all the authors. Y.S.J. supervised the overall study.

## Figure legend

**Extended Data Fig. 1.**
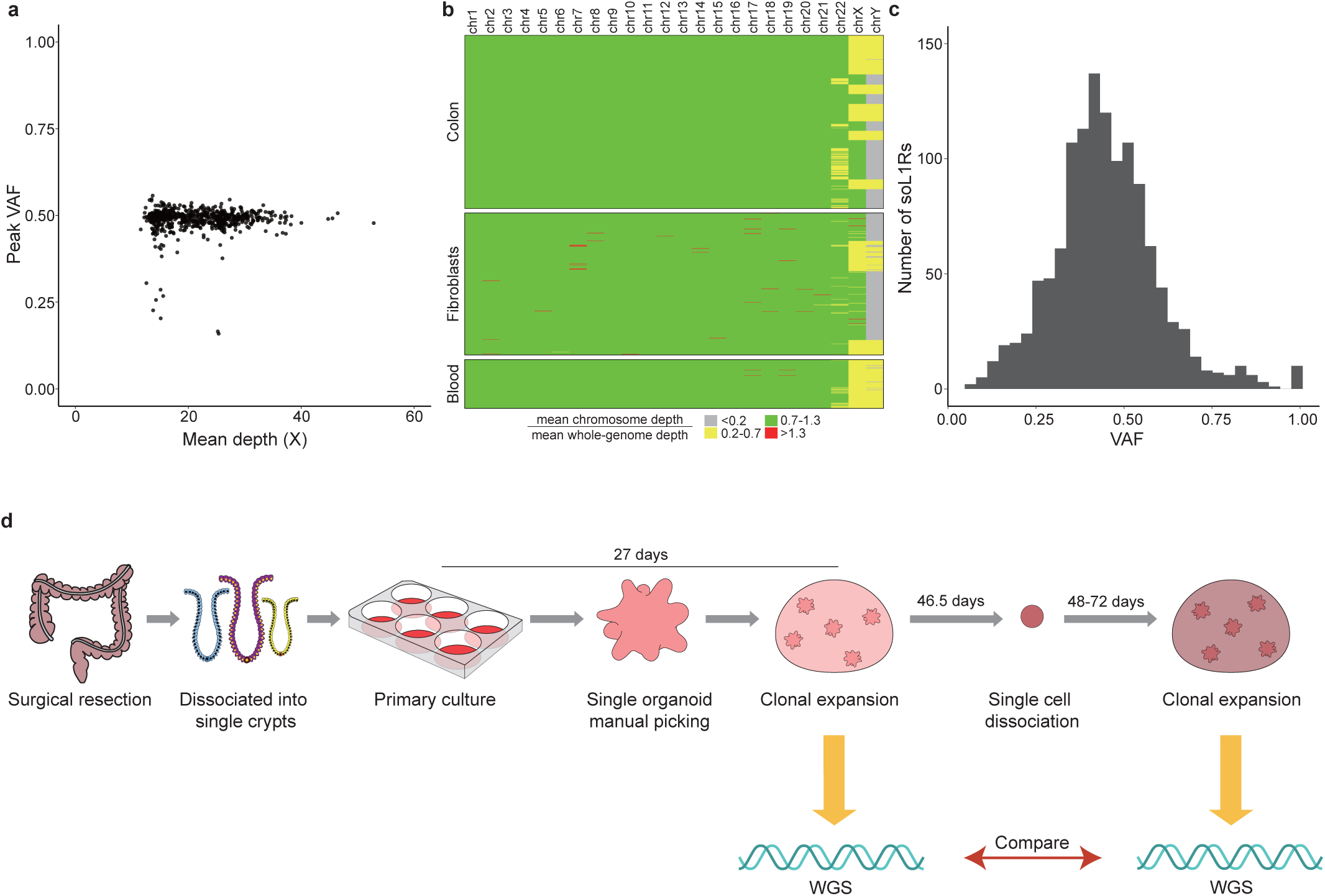
Clones for detection of soL1Rs. **a**, A scatter plot showing mean sequencing coverage of a clone and peak VAF of somatic mutations. Most clones showed their peak VAFs around 0.5, indicating that they were established from a single founder cell. **b**, Chromosome level copy number changes of the 880 clones. No significant genome-wide aneuploidy was detected, supporting genomic stability during clonal expansion of normal single cells. **c**, The distribution of VAFs of 1,236 soL1Rs identified in normal colorectal clones. A peak VAF near 0.5 suggests that the vast majority of soL1Rs detected were shared by all cells in a clone, therefore unlikely to be acquired during cell culture. **d**, Experimental design for estimating the rate of L1 retrotransposition during culture. For whole-genome sequencing of clones, a single crypt from the surgical specimen (naturally clonalized) was cultured for 27 days. The second clonalization was conducted after additional culture of 46.5 days on average (ranging between 43-50d). From 13 pairs of whole-genome sequences of early and late clones, we found ten new clonal soL1R events specifically in late clones, which should be acquired during cell culture before the second clonalization, or ∼73.5 days (27d+46.5d). This allowed us to calculate the culture-associated soL1R rate: 10 soL1Rs / 13 clones / 73.5d = 0.01 per clone per day. Using the rate, we can estimate the upper boundary of the rate, which is 0.01 per clone per day * 27.days = 0.28 soL1Rs, assuming the expansion of the recent common ancestral cell (MRCA) at day 27. Of note, the lower boundary is 0 if the expansion of the MRCA occurred at day 0. It suggests that the proportion of the culture-associated soL1Rs is 9% at maximum, which is 0.28/3.044*100 (3.044 soL1Rs detected per clone, 1,236 soL1Rs from 406 clones).

**Extended Data Fig. 2.**
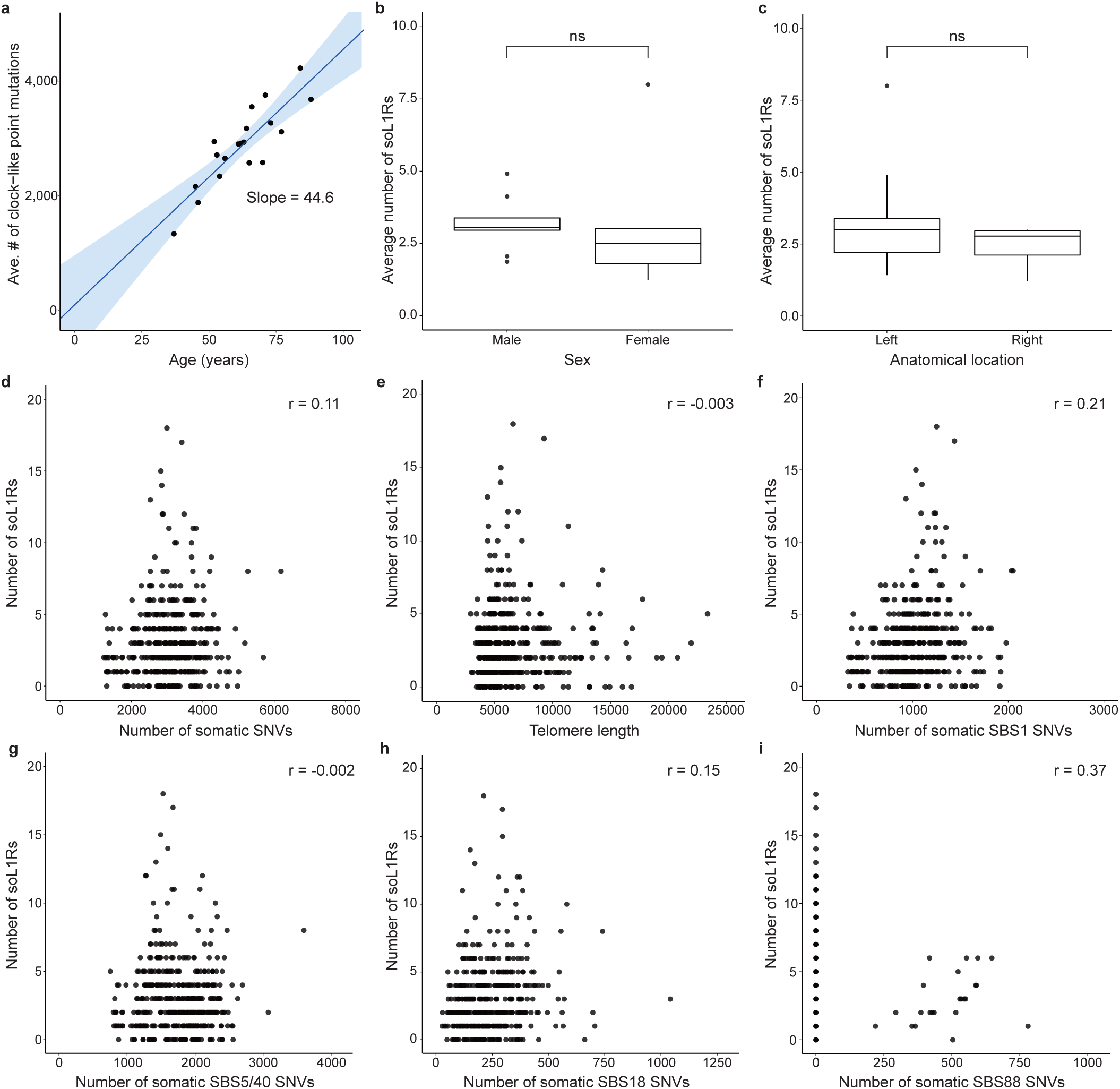
Associations between soL1R burden and other genomic features of clones. **a**, Linear regression between the average number of clock-like point mutations in the colorectal clones and the age of sampling. Blue line represents the regression line (44.6 point mutations per year), and shaded areas indicate its 95% confidence interval. The rate is consistent with the rate previously estimated in the colon (43.6 mutations per year from Lee-Six et al., Ref. 73). **b, c**, Comparison of the average number of soL1Rs per individual across sex (b) and anatomical location of the colorectal crypts (c). **d-i**, Relationship between the number of soL1R for each colorectal clone and the number of somatic SNVs (d), telomere length (e), the number of somatic SBS1 SNVs (f, clock-like mutations by deamination of 5-methyl cytosine), the number of somatic SBS5+SBS40 SNVs (g, clock-like mutations by unknown process), the number of somatic SBS18 SNVs (h, possibly damage by reactive oxygen species), and the number of somatic SBS88 SNVs (i, damage by *pks+ E. coli*). No obvious association was found. ns, not significant; soL1R, somatic L1 retrotransposition.

**Extended Data Fig. 3.**
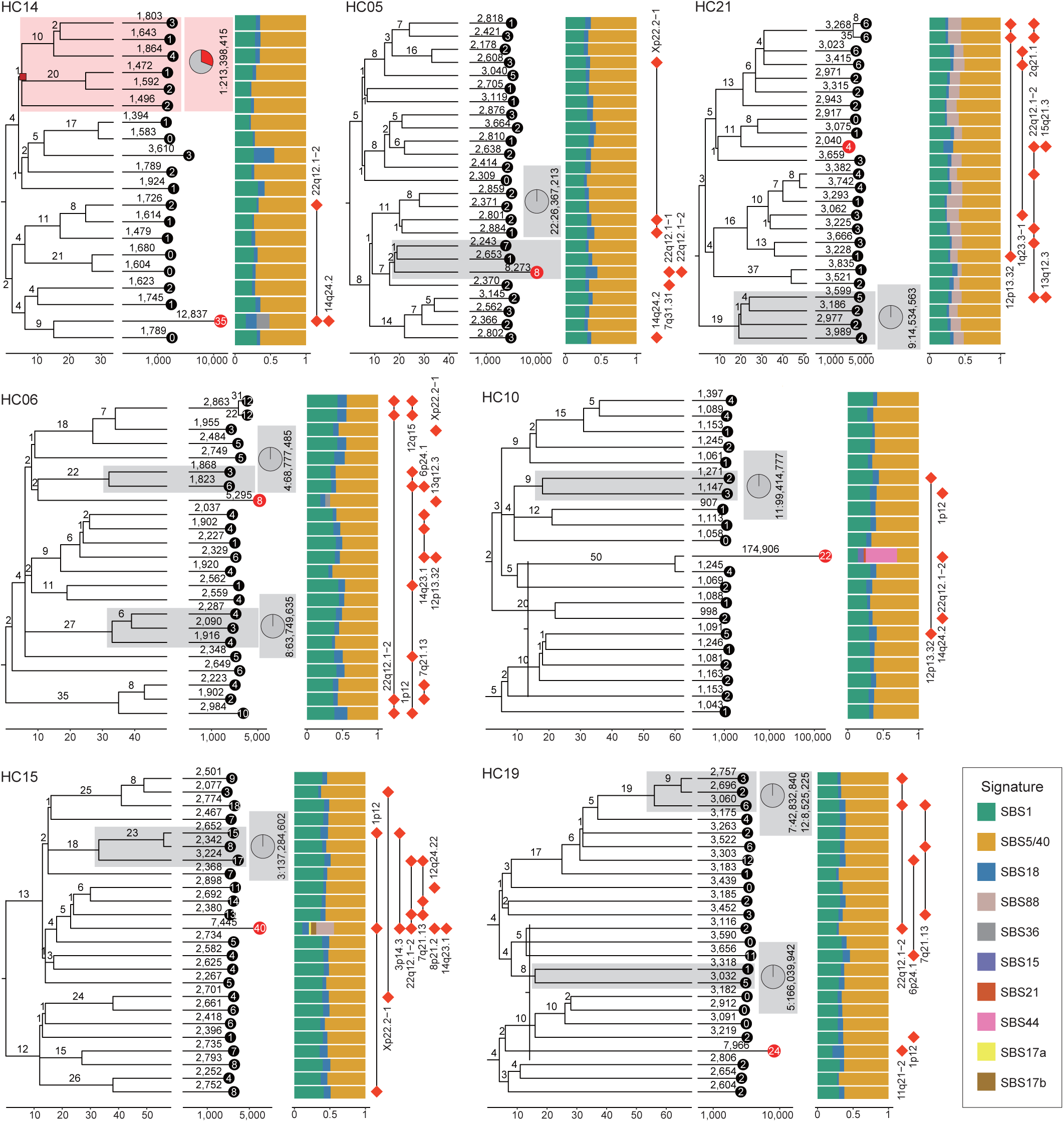
SoL1Rs on the developmental phylogenies of the clones from the seven individuals with early embryonic soL1R events. Early phylogenies of colorectal clones and the matched cancer tissue are shown in seven individuals who have shared soL1Rs among clones. Branch lengths are proportional to the molecular time measured by the number of somatic point mutations. The numbers of branch-specific point mutations are shown with numbers. The filled circles at the ends of branches represent normal clones (black-filled circles) and the cancer clone (red-filled circles). The numbers within the filled circles show the number of soL1Rs detected from the clones. Shaded areas indicate somatic lineages with shared soL1Rs. The genomic location of the shared soL1R insertions and the proportion of the blood cells carrying the soL1Rs are shown by genomic coordinates and pie charts. Coloured bars on the right side represent the proportion of mutational signatures attributable to the somatic point mutations. Orange diamonds show hot L1 sources (origin), which caused transduction events across the colorectal clones.

**Extended Data Fig. 4.**
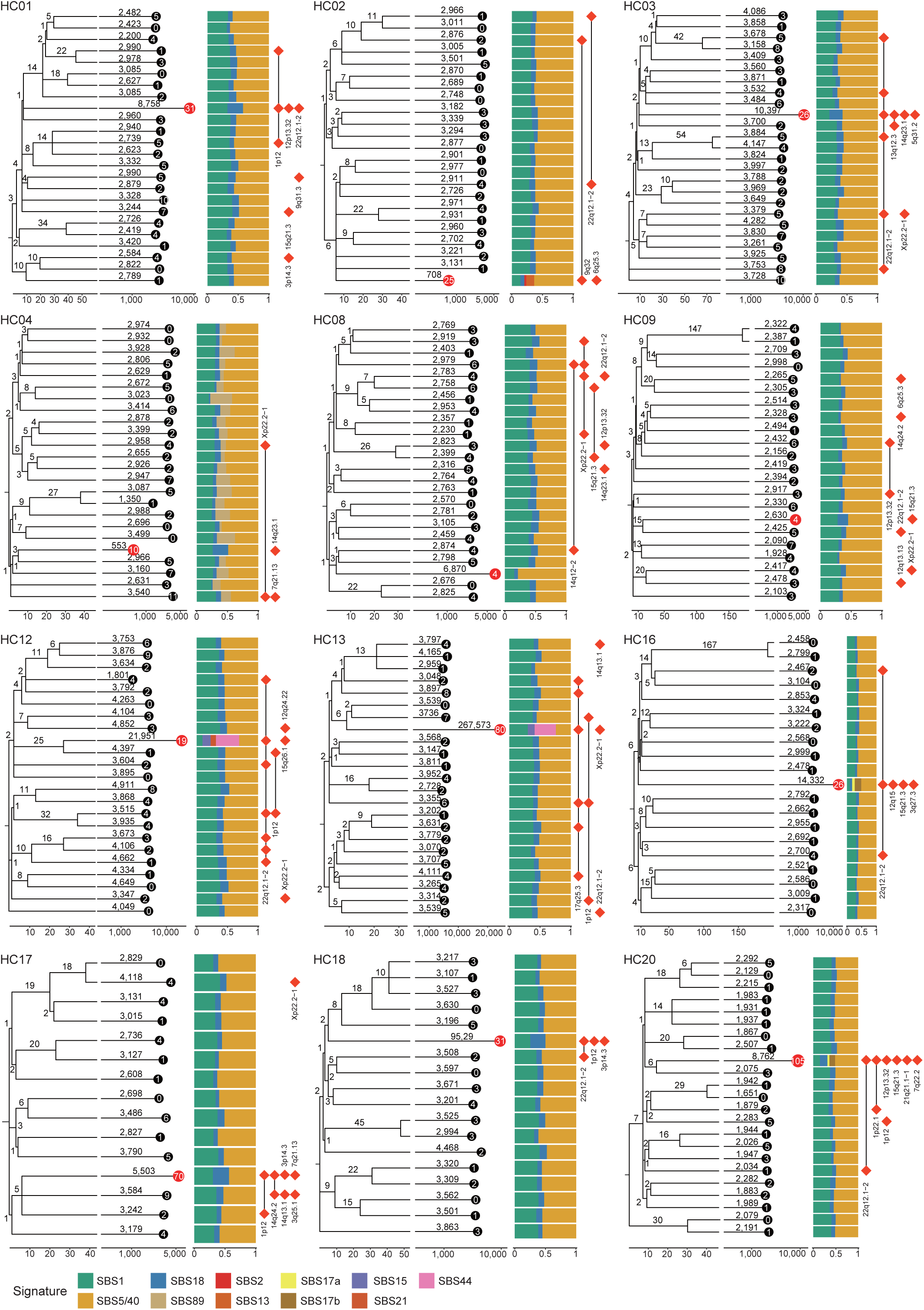
SoL1Rs on the developmental phylogenies of the clones from the 12 individuals without early embryonic soL1R events. Early phylogenies of colorectal clones and the matched cancer tissue are shown in 12 individuals who have no shared soL1Rs among clones. Branch lengths are proportional to the molecular time measured by the number of somatic point mutations. The numbers of branch-specific point mutations are shown with numbers. The filled circles at the ends of branches represent normal clones (black-filled circles) and the cancer clone (red-filled circles). The numbers within the filled circles show the number of soL1Rs detected from the clones. Coloured bars on the right side represent the proportion of mutational signatures attributable to the somatic point mutations. Orange diamonds show hot L1 sources (origin), which caused transduction events across the colorectal clones.

**Extended Data Fig. 5.**
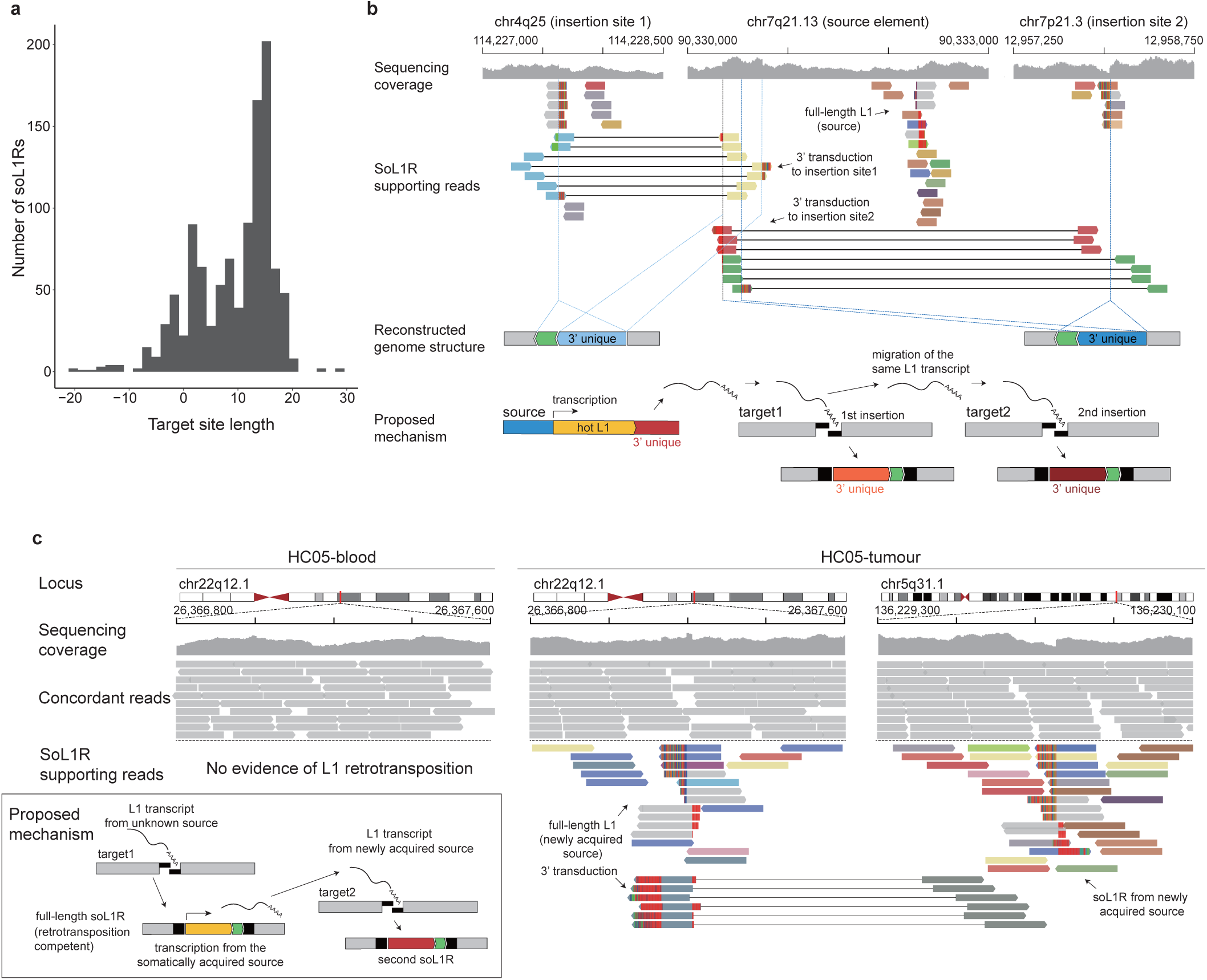
Genomic characteristics of soL1R target regions and examples of soL1Rs providing insights into the L1 dynamics in somatic cells. **a**, The distribution of lengths of target site duplications at insertion points. Positive and negative target site lengths indicate target site duplication and deletion, respectively. **b**, An example of a clone showing evidence of two transduction events from a single L1 transcript. A hot L1 located at 7q21.13 generated orphan transductions at two different target sites (4q25 and 7q21.3) with the same positions of their poly-A tails in the transduced sequences. **c**, An example of a soL1R event (transduction) induced from a somatically acquired L1 source. HC05 tumour has a hot L1 in 22q12.1 (middle) which is not found in the germline of HC05 (blood; left). The 22q12.1 caused a transduction event at 5q31.1 (right) in the tumour, suggesting secondary transduction from the new hot L1 somatically acquired. The proposed order of events is summarized in the lower-left panel.

**Extended Data Fig. 6.**
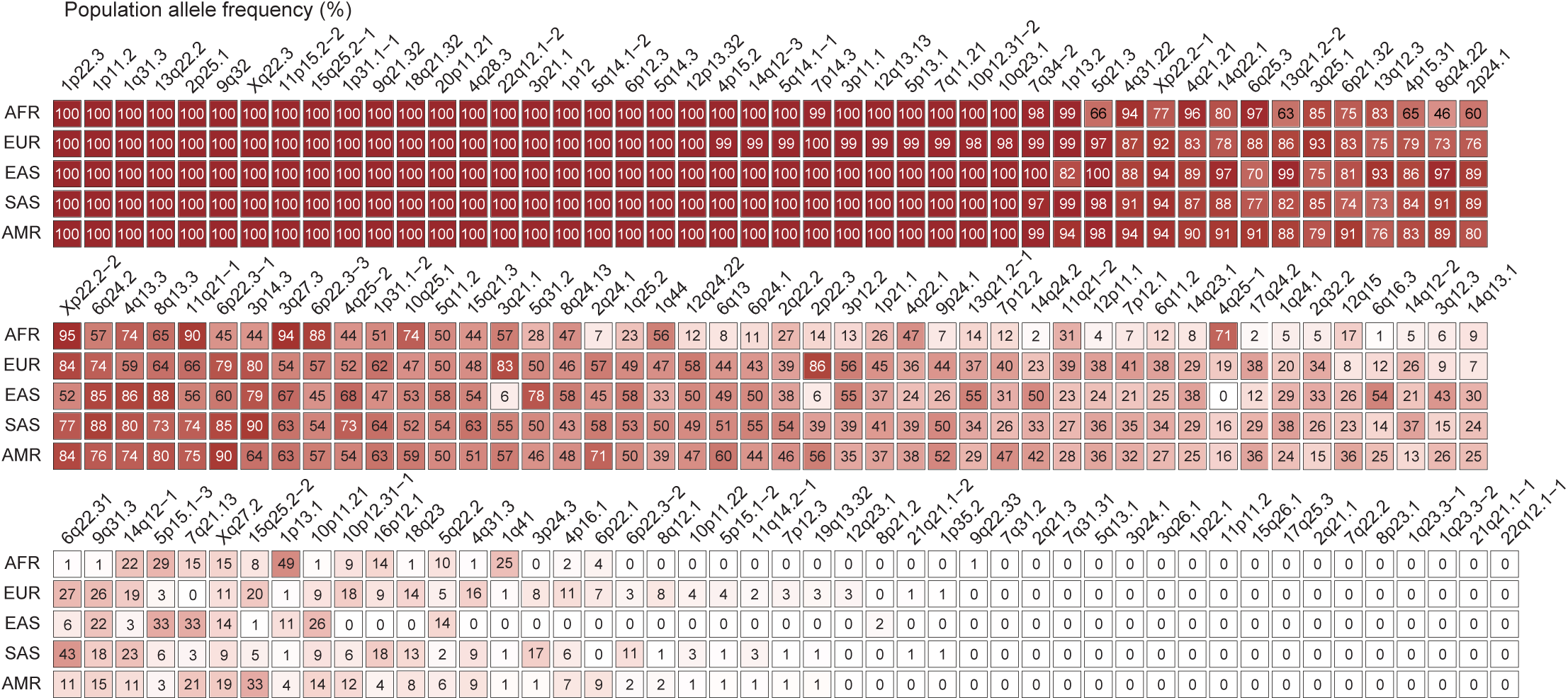
Population allele frequency of 139 hot L1 sources. The population allele frequency of 139 hot L1s (124 previously known + 15 novel sources) was calculated from a panel of 2,860 individuals from 5 ethnic groups. The accurate genomic positions of hot L1s are available in Supplementary Table 3. AFR, African (n=714); EUR, European (n=588); EAS, East Asian (n=646), SAS, South Asian (n=538); AMR, Ad Mixed American (n=374).

**Extended Data Fig. 7.**
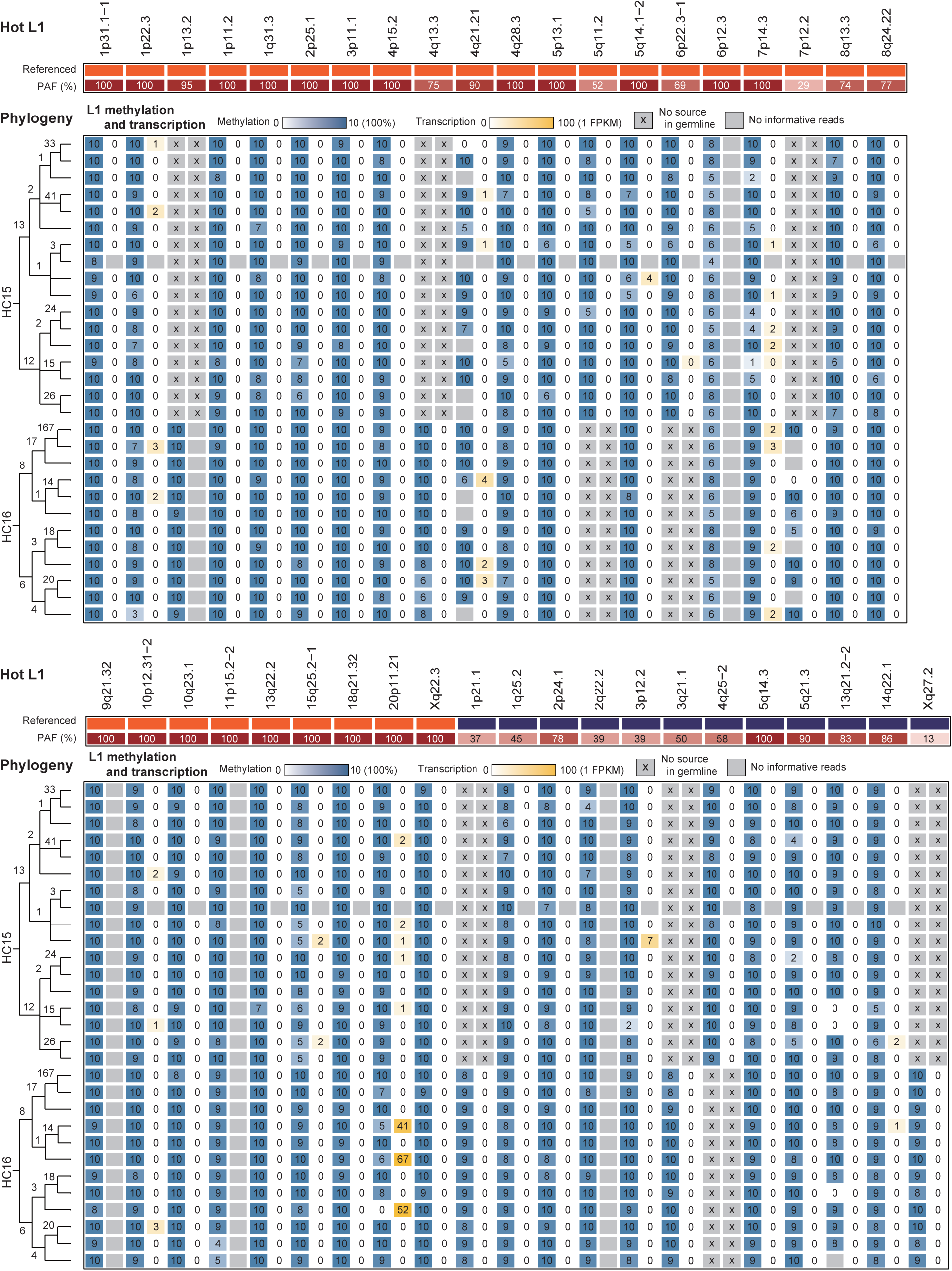
Panorama of DNA methylation status and read-through RNA expression levels. It is an extended version of Fig 4f encompassing an additional 41 hot L1s that are present in the germline of HC15 and HC16. Hot L1s not included in Fig 4f and Extended Data Fig. 7 are not present in the germline of HC15 and HC16. The developmental phylogenies of colorectal clones in HC15 and HC16 are shown on the left side with number of mutations (molecular time). PAF, population allele frequency; FPKM, Fragments per kilobase of transcript per million.

