## Supplementary Discussion for "Extensive mosaicism by somatic L1 retrotransposition in normal human cells"

For the detection of somatic mutations in single cells, a few complementary techniques have been used^1^. In normal intestinal crypts, containing a few thousand cells, a few stem cells continuously compete with each other^2^, resulting in single-clone compartments in the microscopic structure. As in cancer tissues, the cells in a crypt share all somatic mutations acquired in the genome of the founder stem cell. Using specialized technologies such as laser capture microdissection **(LCM)**, these patches can be physically separated, followed by whole-genome sequencing through an enzyme-based library prep technique that allow a low input amount of genomic DNA^3^. In this way, somatic mutations in the normal colorectal epithelium, particularly single base substitutions and indels, were systematically detected and analyzed^4^.

Although LCM is an efficient technique for exploring point mutations at single-cell resolution, we realized that this approach is not adequate for detecting somatic L1 retrotranspositions **(soL1Rs)**. As mentioned in the main manuscript, critical supporting evidence for soL1R detection includes the detection of clustered reads carrying poly-A tails, which are imprints of reverse transcription of transcripts. However, these poly-A carrying reads are technically depleted in the whole-genome sequences prepared through LCM-based technique^4^ compared to those prepared through clonal expansions (**Supplementary Figs. 1a-1c**). The poly-A dropout greatly reduces the sensitivity and precision of soL1R detection in current tools, i.e., MELT^5^, TraFiC-mem^6^, DELLY^7^, and xTea^8^.

The mechanism underlying the poly-A dropout is unclear. We speculate that it stems from the DNA preparation step: the enzymatic fragmentation, used by Lee-Six et al.^4^, may have a bias or be less effective in fragmenting DNA in A (or T) homopolymeric regions **(Supplementary Fig 2)**. Whichever the reason, we conclude that current LCM-based whole-genome sequencing datasets are practically not suitable for soL1R analysis until substantial improvements in the library preparation step, bioinformatics tools, or both.

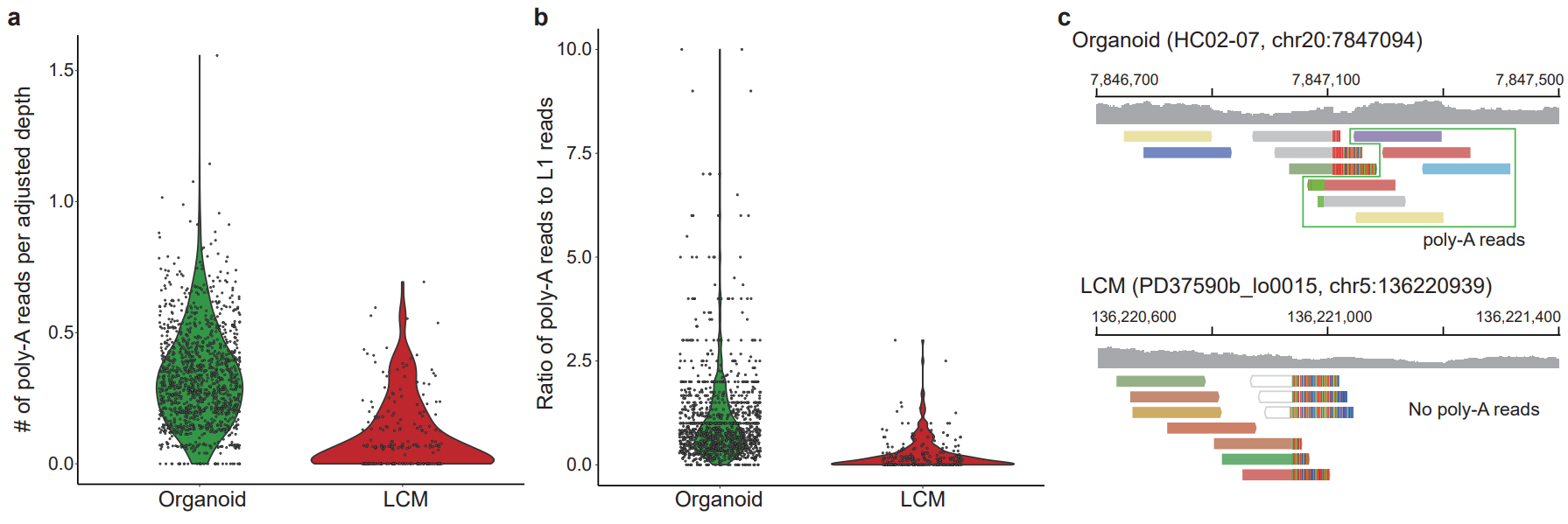

**Supplementary Fig. 1. Lack of poly-A reads in whole-genome sequences prepared using LCM**

**a**, Number of poly-A reads per adjusted depth. The soL1Rs detected in whole-genome sequences prepared using LCM have fewer poly-A reads per depth compared to those from clonal expansion (0.33 vs. 0.11 on average for clonal expansion and LCM, respectively, *P*=3.1×10^-59^, two-sided t-test). Poly-A reads indicate the read pairs with poly-A sequences in the reads or their mate reads. Sequencing depth was adjusted with purity, which is estimated as two times peak VAF. soL1R, somatic L1 retrotransposition; LCM, laser capture microdissection; VAF, variant allele fraction. **b**, Ratio of poly-A reads to L1 reads. The ratio is lower in soL1Rs detected in whole-genome sequences prepared from LCM than those from clonal expansion (1.07 vs. 0.26 on average for clonal expansion and LCM, respectively, P=2.2×10^-72^, two-sided t-test). L1 reads indicate the read pairs with L1 consensus sequences in the reads or their mate reads. **c**, Examples of soL1Rs detected in whole-genome sequences prepared using clonal expansion (top) and LCM (down).

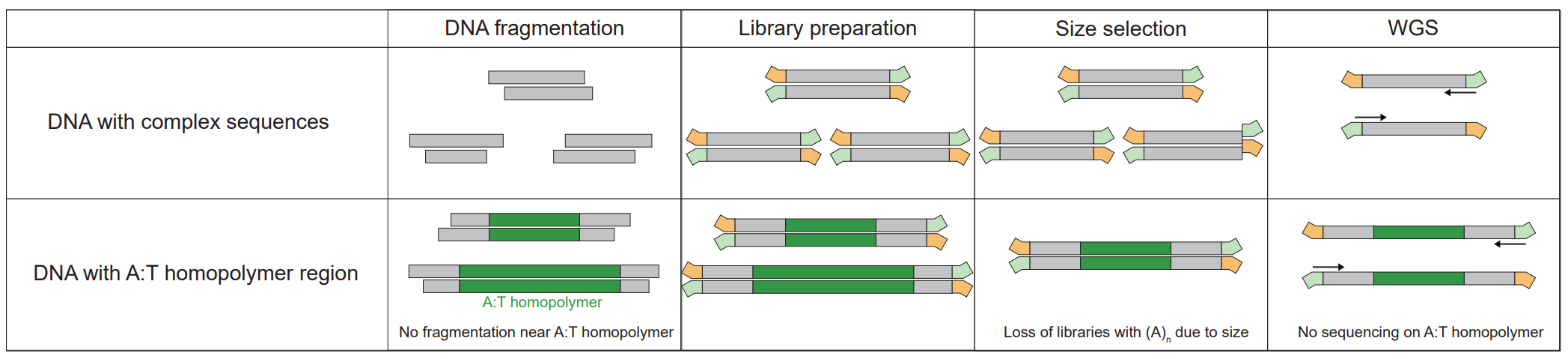

**Supplementary Fig. 2. Proposed mechanism underlying poly-A dropout in whole-genome sequences prepared using LCM.**

If DNA with A:T homopolymer regions are less vulnerable to enzymatic fragmentation, A:T homopolymer regions are likely to be included in large-sized libraries, which could be eliminated during the size selection process. Even if A:T homopolyer containing libraries survive, A:T homopolymer region far from the adapter at both ends would not be sequenced. LCM, laser capture microdissection.
